# Loss of function from widely distributed, synonymous mutations at single codons

**DOI:** 10.1101/684092

**Authors:** Kritika Gupta, Jyothi Prabha, Soumyanetra Chandra, Shruti Khare, Sonali Vishwa Mohan, Pehu Kohli, Aparna Asok, Harsha Gowda, Arati Ramesh, Raghavan Varadarajan

## Abstract

Mutational tolerance inferred from laboratory-based mutational studies is typically much higher than observed natural sequence variation. Using saturation mutagenesis, we show that the *ccdA* antitoxin component of the *ccdAB* toxin-antitoxin system is unusually sensitive to mutation with over 60% of mutations leading to loss of function. Multi-base synonymous mutations at a codon display enhanced propensity to show altered phenotypes, relative to single-base ones. Such mutations modulate RNA structure, leading to altered relative translation efficiencies of the two genes in the operon, and a CcdA:CcdB protein ratio below one. These insights were used to predict and experimentally validate synonymous mutations that lead to loss of function in the unrelated *relBE* operon as well as the *lacZ* gene. Thus, synonymous mutations can have significant phenotypic effects, in the absence of overexpression or extraneous reporters. More generally, proteins are likely more sensitive to mutation than inferred from previous saturation mutagenesis studies.

## Introduction

The three-dimensional structure of a protein is dictated by its amino acid sequence and in most cases the structure determines protein function. Classically, mutational phenotypes are used to establish gene function and identify residue specific contributions to protein function. Non-synonymous and nonsense mutations are often selected against as they lead to partial or complete loss of protein function. Synonymous mutations do not change the amino acid sequence and are expected to be neutral. It is now known that synonymous mutations can in some instances be under selection pressure, but this is believed to be relatively rare (1). Synonymous mutations have been reported to be significantly associated with over 50 human diseases (2) emphasizing need for further study of such mutations.

To decipher the molecular basis of the effects of synonymous substitutions, many studies have been performed, both at the single gene and whole genome level. Some of the mechanisms known to be involved in phenotypes observed upon synonymous mutation are; direct effect on protein expression and folding (3), effects on mRNA stability due to change in secondary structure or mRNA decay (4) changes in protein expression, or bias due to tRNA abundance (5) and effects on translation initiation (6), elongation or termination (3). Synonymous mutations were previously thought to be silent probably due to limited sensitivity and throughput of the assays and proteins used in traditional studies. Mutational sensitivities in experimental studies were generally found to be quite low, leading to the expectation that large sequence variation should exist among natural homologs of the protein, but this was typically not found (7). Following the advent of high-throughput mutagenesis studies coupled to deep sequencing, most studies showed little or no effect of synonymous mutations, with a few exceptions (8, 9). One limitation in these studies is the use of artificial stress conditions, such as antibiotic stress, that complicates direct evolutionary interpretation (10). Most such studies also employed heterologous proteins over-expressed in *E. coli* or other model organisms, limiting the ability to understand the intrinsic *in vivo* determinants of such phenotypes, and the effect of synonymous mutations in a native context on fitness. Low or moderately expressed genes are likely not affected by codon bias as much as highly expressed genes, as the abundance of cognate tRNA may not be a limiting factor for these genes. Under physiological conditions, especially in cases where the gene expression is low, it therefore becomes important to study the contributions of individual causal factors to variation in gene expression.

It is also known that there is a selection for local RNA secondary structure at a genome-wide level (11) and that reduced local RNA structure and not codon rarity contributes to increase in expression levels, especially at the N-terminus of genes (12). With the availability of whole genome transcriptomic, proteomic as well as ribosome profiling data, the determinants of efficient transcription, translation as well as efficient resource allocation in bacteria are slowly beginning to be understood. One way to study the effects of mutations on mRNA translation, independent of mRNA stability, is to monitor the changes in levels of proteins expressed from the same polycistronic mRNA. This assumes that any global effects would alter levels of all genes in the operon. Given a common mRNA sequence, one of the important ways in which modulation of relative expression of the genes in an operon can be controlled, is through differential translation, possibly driven by distinct structures in mRNA in adjacent genes (13). Differential transcription or differential mRNA stability is unlikely in the case of genes encoded in polycistronic mRNAs except in cases where there could be internal transcription initiation or termination sites. For genes in an operon to display differential rates of protein production, internal initiation of translation is likely essential.

Toxin-antitoxin (TA) systems in bacteria are an attractive model to study co-regulation of gene expression as well as transcriptional and translational coupling (14). For most type II TA systems, the antitoxin and the toxin genes are expressed as part of a single operon (15). Although the distance between the stop codon of the first gene and start codon of the second gene can vary, bioinformatics studies have shown that in the case of TA systems it is between 0-5 nucleotides (15). In some cases, the two ORFs even overlap with each other (16). Post-translationally, the two proteins are subjected to differential degradation. The ratio of antitoxin to toxin is expected to diminish during stress conditions, as in the case of persisters, where it is thought that the accumulation of the toxin leads to metabolic dormancy by slowing vital cell processes such as replication, transcription and translation (17). Although conditional co-operativity is known to be a dominant mechanism in regulating the transcription of most TA systems (18), mechanisms that can lead to differential levels of the toxin and the antitoxin protein from a common mRNA are not well understood. For conditional co-operativity to manifest, the default assumption is that the antitoxin is present at significantly higher levels than the toxin, a notion which has not been rigorously tested previously. Given the higher rate of degradation for the antitoxin due to its disordered nature, the antitoxin needs to be produced at a higher rate than the toxin, for the cells to be viable. While levels can be in principle regulated at both the transcriptional as well as the translational level, RNA” seq data from *E.coli* indicate that mRNA levels for the toxin and the antitoxin are not significantly different, as is expected for genes encoded by a single polycistronic mRNA (16). Ribo-seq data available for TA systems indicate that the protein synthesis rates (calculated from the ribosome densities) for the antitoxin are at least 2 fold higher than the toxin, for six TA pairs for which sufficient data is available (16). These observations together indicate that there are indeed nucleotide sequence dependent features, which allow regulation at the level of translation, leading to differential synthesis rates for the two genes in the operon. We therefore investigated the molecular bases for such regulation by carrying out saturation mutagenesis studies of the antitoxin *ccdA*, from the CcdAB TA system in *E. coli*.

The *ccd* operon encodes a labile 72 residue CcdA antitoxin, that prevents killing of *E. coli* cells by binding to the 101-residue toxin, CcdB. Both genes are co-expressed in low amounts in F-plasmid bearing *E. coli* cells, and their expression is autoregulated at the level of transcription (19). If the cell loses F-plasmid, the labile CcdA is degraded by the ATP dependent Lon protease, releasing CcdB from the complex to act on its target DNA gyrase, which eventually leads to cell death (20). Therefore, mutations which disrupt CcdA antitoxin function lead to cell death, under normal conditions. Using saturation mutagenesis, we find that the labile antitoxin, CcdA apparently shows considerably higher mutational sensitivity than most proteins studied to date (10). Several synonymous point mutations in CcdA lead to loss of function in a manner uncorrelated to codon preference. Our data suggests that changes in mRNA structure upon mutation result in lowering of the CcdA: CcdB ratio *in vivo*, through a self-amplifying feedback loop, ultimately leading to cell death. The *ccd*, and likely most other TA operons, are highly sensitive genetic circuits that can be used to probe effects of mutations on gene function, *in vivo*. Most studies till date suggest that multiple synonymous mutations are required for a discernible phenotype. Subtle differences in fitness that are often undetectable in laboratory experiments can become fixed in the population over long timescales (7, 21) and laboratory selection conditions can differ greatly from those experienced during natural selection. Hence, to properly understand the contribution of single mutations, it is useful to have experimental systems, such as the present one, where small changes in protein activity have substantial phenotypic effects that are largely independent of external conditions.

## Results

### Unexpectedly high mutational sensitivity in the *ccdA* antitoxin gene encoding a natively unfolded protein

The *ccd* operon encodes for a labile 72 residue CcdA antitoxin, that prevents killing of *E. coli* cells by binding to the 101-residue toxin, CcdB. Mutations which disrupt CcdA antitoxin function lead to cell death. The ccd operon consisting of the promoter, ccdA and ccdB genes was cloned into the pUC57 plasmid (Supplementary Figure 6). A site-saturation mutagenesis (SSM) library of CcdA was prepared using NNK primers (22). The pooled mutant library of CcdA, transformed in sensitive and resistant strains (see Materials and Methods), was subjected to deep sequencing to estimate the relative ratio of the mutants in selected vs. unselected conditions (Supplementary Figure 1). Out of a possible 2272 (71 positions*32 codons) mutants expected by NNK codon mutagenesis of the CcdA gene, reads for ~1700 mutants were available in the strain resistant to the toxin activity (represents the unselected library), with an average of ~300 reads per mutant. Only mutants having more than five reads in the unselected conditions were further analyzed. Deep sequencing was performed on two biological replicates, with good agreement between common mutants. Relative sensitivities of the total pooled library, mutants in the DNA binding domain, and in the CcdB binding domain were also assessed *in vivo* (Figure 1). A drastic reduction of cell growth in the sensitive strain shows that most CcdA mutants in the library have an inactive phenotype. We defined depletion ratio as the ratio of the normalized reads in the sensitive strain to the normalized reads in the resistant strain, for a given mutant. The distribution of depletion ratios amongst the CcdA mutants was assessed by plotting the frequency of mutants as a function of increasing depletion ratio. The distribution was centered on a peak at depletion ratio of 0.3 (Figure 1). ~60% of mutants had depletion ratio <0.3. The phenotype for a subset of these mutants was tested in the sensitive strain, by individual transformations and plating, and confirms that mutants with a depletion ratio <0.3 showed significant growth defects in the sensitive strain (Supplementary Figure 1). Overall, our saturation mutagenesis studies indicated that in an operonic context (Supplementary Figure 6), the antitoxin, CcdA has an exceptionally high mutational sensitivity, with ~60% of the mutants exhibiting an inactive phenotype and ~83% showing decreased activity relative to WT.

**Figure 1:**
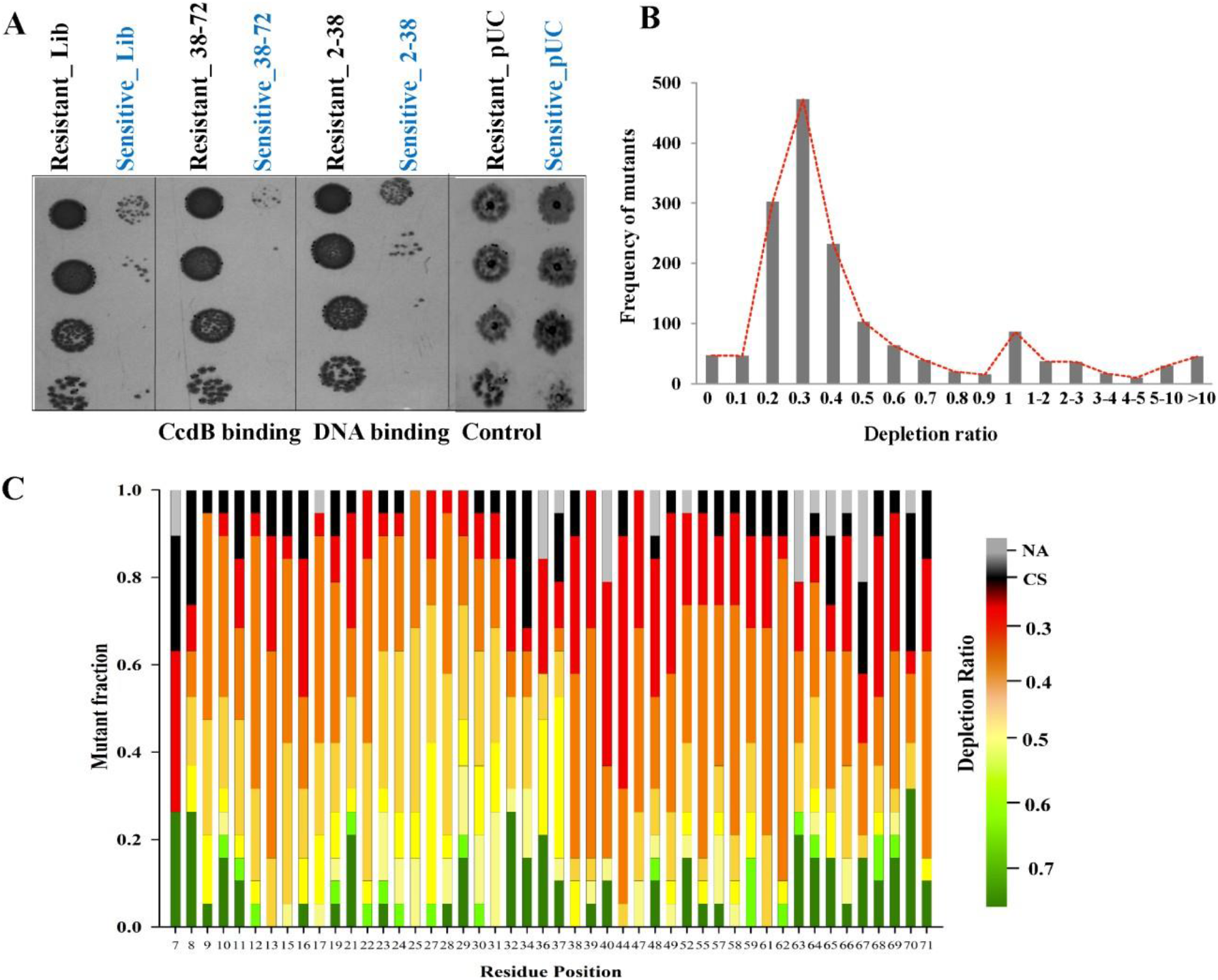
High mutational sensitivity of ccdA antitoxin gene in native operon context in *E. coli*. (A) Relative mutational sensitivities of the various ccdA mutant libraries. Pooled libraries of all ccdA mutants, mutants in the DNA binding domain, and in the CcdB binding domain were transformed individually in the resistant and sensitive strains of *E. coli*. The transformation mix was spotted on LBa*mp* agar plates in two-fold serial dilutions. The drastic reduction of cell growth in the sensitive strain shows that several ccdA mutants in the library have a loss of function phenotype. (B) Distribution of the depletion ratios for the ccdA mutants in the library inferred through deep sequencing data. For each mutant in the library, ratio of the normalized reads in the sensitive strain to the normalized reads in the resistant strain was calculated. A low depletion ratio (<0.3) is indicative of low mutant activity. All mutants having less than five reads in the resistant strain were not considered for analysis. (C) Phenotypic landscape for different residues in CcdA inferred from the deep sequencing data. For each of the residue positions in CcdA, the fraction of mutants having a given depletion ratio (represented in different colors) is plotted. Black color represents fraction of mutants having codon specific (CS) phenotype and grey represents mutants not available. A large number of mutants have a loss of function phenotype, the fraction being higher for the C-terminal residues 37-72. Only positions which have more than 13 mutants available have been plotted here.

### Synonymous inactive mutations are unequally distributed throughout the *ccdA* gene

Surprisingly, several mutants at multiple positions display codon specific activity effects (Figure 1). For example, for the D71R mutation, two of the mutant codons CGG and CGT show an active phenotype, whereas the AGG codon shows an inactive phenotype (Appendix Table 1). Interestingly, about 90% of all inactive mutants in the library have two or three base mutations, indicating a crucial role of the nucleotide sequence, rather than just the protein sequence in determining the mutant phenotype. At many of the positions even the WT synonymous codons were found to be inactive (Figure 2). From a total of 71 synonymous mutants available in the library, 38% of the mutations had a depletion ratio less than 0.3, including 16 single base substitutions. All but one, three-base substitution among the synonymous mutations were inactive. Phenotypes have been confirmed, in several cases by constructing individual mutants and screening for activity on plates (Supplementary Figure 2A). The distribution of all available synonymous mutations along the length of the *ccdA* gene indicated an increased frequency of synonymous inactive mutations from residues 8-31 and 63-71 (Figure 2B). Such a trend is also visible for positions manifesting codon specific activity (Figure 2C).

**Figure 2:**
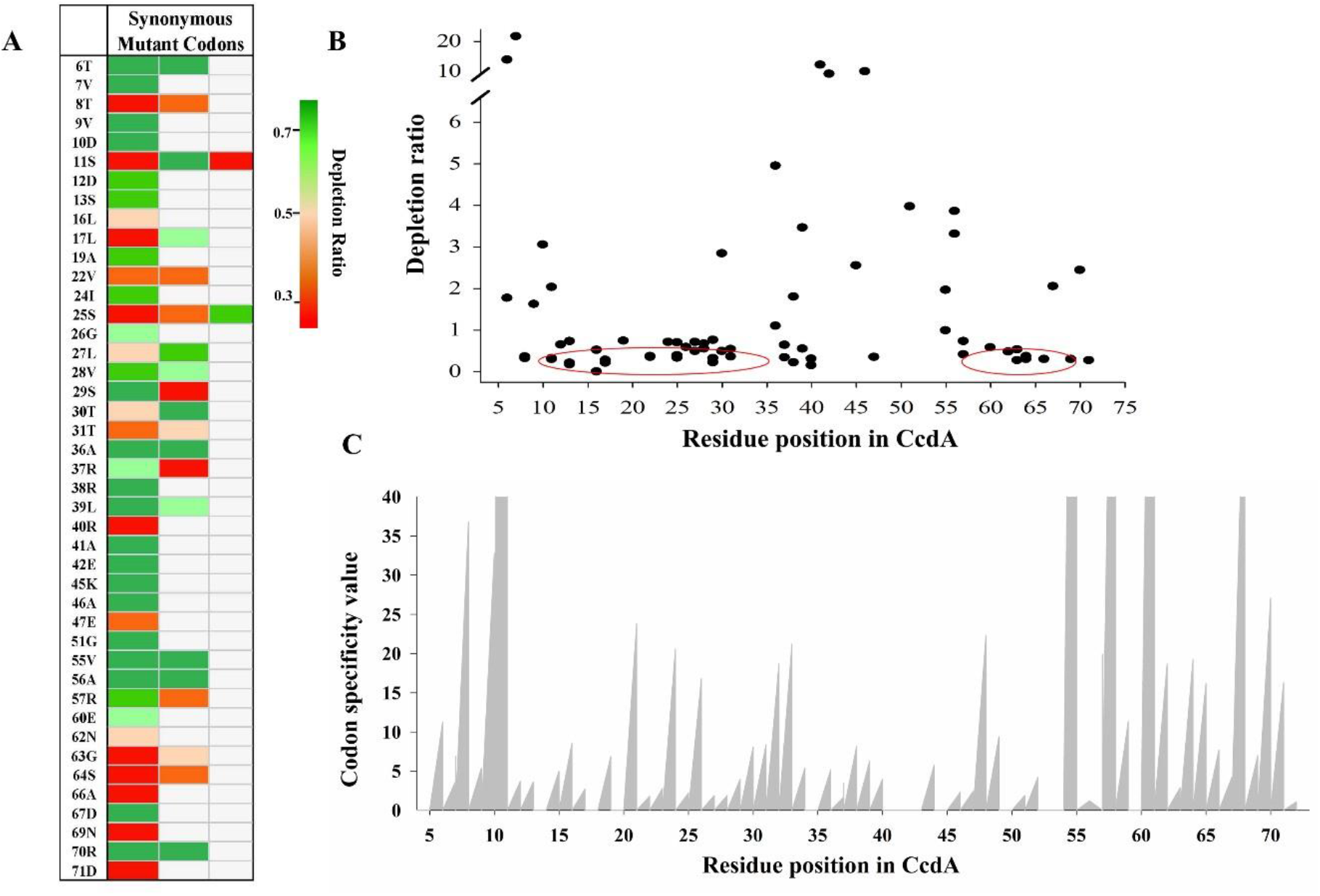
Synonymous inactive mutations are unequally distributed throughout the *ccdA* gene. (A) Mutational sensitivities of all available WT synonymous mutant codons in the CcdA library. Mutant activity has been colored from green to red in order of decreasing depletion ratio. (B) Distribution of depletion ratios for single synonymous mutants with respect to the WT has been plotted as a function of residue position. Clusters of loss of function mutations are marked in red circles. (C) Distribution of the codon specific effects as a function of residue position in CcdA. Codon specific value for each amino acid substitution was calculated by dividing the maximum depletion ratio for a codon by the minimum depletion ratio for a codon, coding for the same amino acid.

### Strength of CcdB ribosome binding site correlates moderately with CcdA mutant phenotype

A primary sequence element that is known to be a strong determinant of the efficiency of translation coupling is the Shine Dalgarno sequence. Early studies by Das and Yanofsky on the Trp operon clearly implied that translation initiation at any start site located near a functional stop codon may be influenced by the strength, sequence and location of the SD region and its spacing from the start codon (23). The Ribosome Binding Site (RBS) for the ccdB gene lies within the ccdA coding region, primarily between the 70^th^ and 71^st^ amino acids. Therefore, mutations in these residues not only may affect CcdA-CcdB binding but may also affect the relative CcdA and CcdB levels by changing the strength of the RBS for CcdB. An analysis of all available codon substitutions at the 70^th^ and 71^st^ residue of CcdA from the deep sequencing data indicated a modest correlation between the strength of the RBS for CcdB expression and the depletion ratio of these mutants (Figure 3A). This indicates that modulation in the strength of the CcdB RBS through mutations in the *ccdA* sequence is an important determinant of mutant phenotype. The molecular basis of one such mutation D71R was probed. Since two of the arginine codon mutants had an active phenotype, it is unlikely that the inactive phenotype observed for the AGG codon is due to effects on protein stability or protein-protein interaction. It is known that the consensus sequence for the RBS, having the maximal strength is 5’ AGGAGG 3’ (based on the corresponding complementary sequence of 16S rRNA, 5’ CCUCCU 3’). Therefore, it is likely that a change from the suboptimal RBS for CcdB present in the *ccdAB* operon, to a consensus RBS, increases the CcdB: CcdA ratio in the cell, thereby leading to cell death. To confirm this, the CcdAB operon having the D71R_AGG mutation in the CcdA gene was cloned under the pBAD promoter in the pBAD24 vector and relative levels of CcdA and CcdB were monitored in the *E. coli* CcdB resistant strain, Top10GyrA. We found that relative to the WT CcdAB construct, the D71R_AGG mutant indeed had higher levels of CcdB protein in the cell (Supplementary Figure 3).

**Figure 3:**
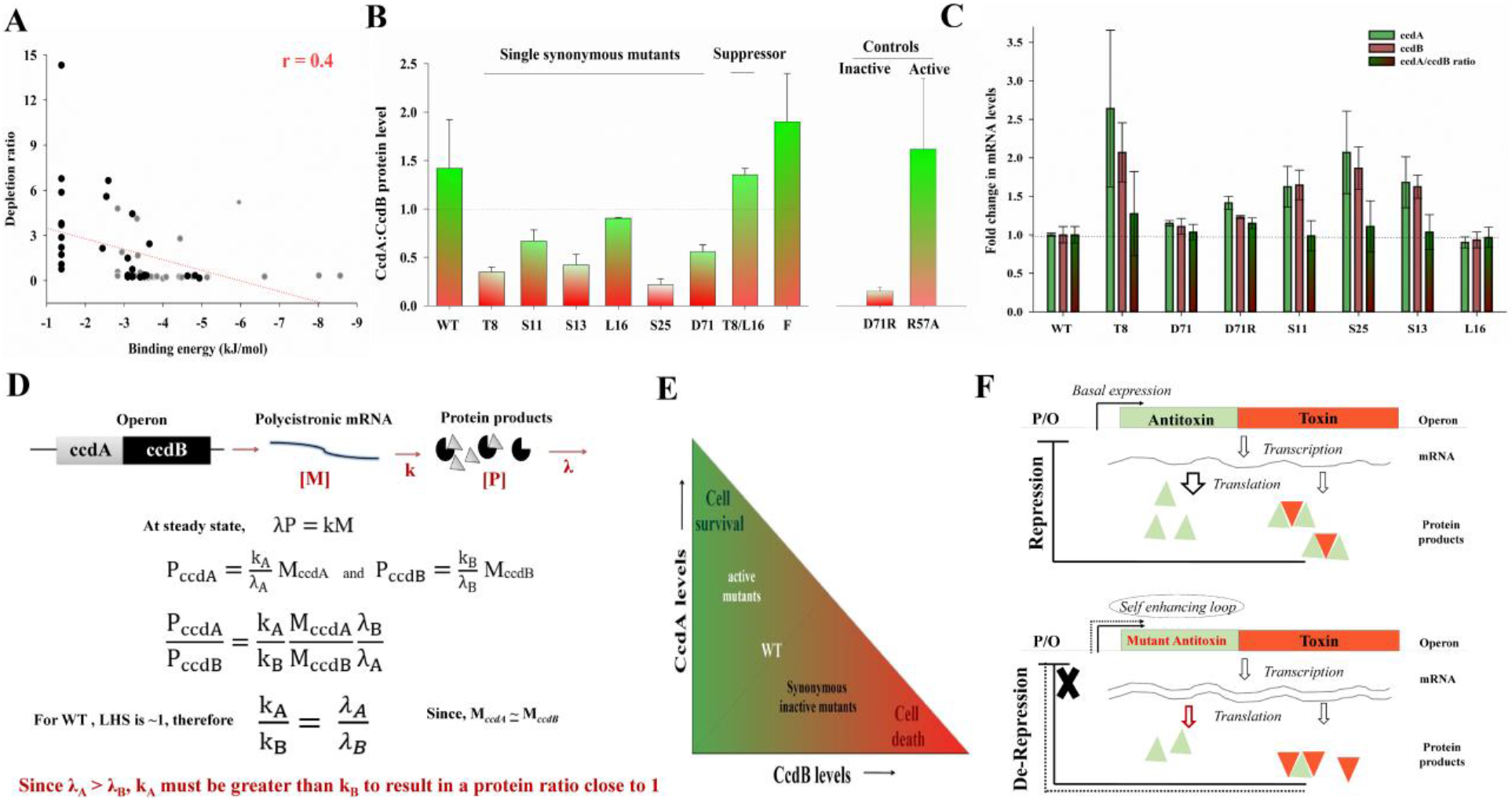
Decreased CcdA: CcdB protein ratio leads to loss of function phenotype of CcdA mutants. (A) Strength of CcdB ribosome binding site correlates moderately with CcdA mutant phenotype. Binding energy to the anti-Shine Dalgarno sequence, for all codon mutants at the 70^th^ (shown in black circles) and 71^st^ residue (in grey circles) positions in CcdA were calculated using RNAsubopt program (Vienna package). This binding energy is measure of the CcdB RBS strength and thereby CcdB protein expression (B) Relative levels of CcdA and CcdB peptides in *E. coli* lysates were determined for WT as well as the single synonymous mutants of CcdA using a quantitative proteomics approach. Most synonymous inactive mutants have decreased CcdA: CcdB protein ratio. ‘F’ indicates the protein ratio in XL-1 Blue strain that bears the ccd operon on WT F-plasmid (C) Changes in relative protein levels cannot be explained by changes in the mRNA levels. mRNA levels of ccdA and ccdB specific transcripts were determined using q-RTPCR. Mean Ct represents the threshold cycle for amplification obtained from duplicate samples from two different experiments. Fold change in ccdA and ccdB transcript levels are with respect to the WT levels. Error bars indicate estimated SD of the measurements. (D) Differential translational efficiency of *ccdA* and *ccdB* genes in the operon. The steady state levels of CcdA (P_ccdA_) and CcdB (P_ccdB_) proteins are assumed to scale linearly as a function of their mRNA concentration (M_ccdA_ and M_ccdB_). [M] and [P] represents mRNA and protein concentrations respectively. Translation efficiency (k_A_ for ccdA and k_B_ for ccdB) is the rate of protein production per mRNA. λ_A_ and λ_B_ are the protein degradation rate constants of CcdA and CcdB. (E) Balance between the steady state *in vivo* levels of the antitoxin CcdA (green) and toxin, CcdB (red). Synonymous mutations in CcdA that lead to loss of activity in the operon context have CcdA: CcdB ratio lower than one, thereby resulting in cell death. A WT CcdA: CcdB ratio of ~1 allows small perturbations in growth conditions to alter the ratio, and switch between phenotypes of cell death and growth as well as facilitating bacterial persistence. (F) Differential translational efficiency of two genes expressed from a polycistronic mRNA affects the phenotype. Mutations in the antitoxin gene can affect translational efficiency thereby leading to changes in the relative steady levels of the protein products in the mutants. As the ratio of toxin: antitoxin exceeds one, the operon is de-repressed leading to an increase in transcription, further enhancing the effect of altered translation efficiencies on relative steady levels of the protein products, leading to cell death.

### Growth kinetics of single synonymous mutants upon overexpression

The above results indicate that synonymous mutant phenotypes in a natural context may differ from non-physiological over-expressed conditions where a strong influence of codon rarity is typically observed (24). To compare the growth of single synonymous CcdA mutants upon overexpression, the WT ccdAB genes, as well as the mutants were expressed from the pBAD promoter under inducing (0.2% arabinose) and repressing conditions (0.2% repressor). The overall growth of the mutants was very similar in the resistant strain, but their growth rates differed in the sensitive strain (Supplementary Figure 2B). Most mutants did not show the drastic growth inhibition that was seen when expressed from their native promoter, but growth rates were relatively lower when expression was repressed. The D71R mutant inhibits cell growth under both repressed and induced conditions, likely by increasing the strength of the RBS for toxin expression. It is probable that increase in inducer leads to an increase in the mRNA: ribosome ratio, such that fewer transcripts will now exhibit a polysomic nature, thereby enhancing coupled translation and maintaining a WT like CcdA:CcdB ratio greater than one. These findings reiterate the fact that mutational effects may be quite different for over-expressed and endogenous genes.

### Decreased CcdA: CcdB protein ratio explains the inactive phenotype of CcdA synonymous mutants

Since the CcdAB operon is repressed under physiological conditions, these proteins are expressed at very low levels and are not detectable on SDS-PAGE. For stable proteins, ribosome density along its corresponding mRNA may provide an estimate of the absolute protein levels in the cells (25). However, for proteins that are subjected to regulated degradation, the determination of absolute levels becomes non-trivial. To monitor if the synonymous inactive mutants exhibit a perturbed CcdA: CcdB ratio *in vivo*, we used a quantitative proteomics approach. Absolute quantitation of endogenous levels of CcdA and CcdB proteins in *E. coli* lysates was done using labelled, synthetic CcdA and CcdB peptides. Lysates from several synonymous point mutants of CcdA that lead to loss of activity in the operon context were subjected to quantitative mass spectrometry analysis.

Although, it is intuitive that the CcdA antitoxin must be present in excess of the CcdB toxin *in vivo* to inhibit toxin mediated cell death, for most TA systems, absolute or relative antitoxin and toxin protein levels are not known. We find that the ratio of the WT toxin and antitoxin proteins in the cell for this CcdAB construct are close to one (Figure 3B) i.e., CcdA and CcdB proteins are present in nearly equal copies in the cell at steady state. For the ccd operon of F plasmid, levels are marginally higher. Synonymous point mutations in CcdA that lead to loss of activity in the operon context have a CcdA: CcdB ratio lower than one, thereby resulting in cell death. For mutants such as the T8 synonymous mutant, this ratio is quite low, whereas for an active mutant, R57A, the ratio is well above one.

The conditional co-operativity model for auto-regulation in toxin-antitoxin systems suggests that an excess of toxin can lead to operon de-repression, as the extended toxin-antitoxin complex is unable to form/bind to the corresponding operator sites on the DNA. At steady state, single synonymous mutants of CcdA that have a lower CcdA: CcdB ratio than WT, are anticipated to cause de-repression of the operon. The degradation rate of the CcdA antitoxin is estimated to be at least five times higher than the cognate toxin CcdB from pulse-chase experiments (26). Therefore, to attain the observed steady state CcdA: CcdB ratio of ~1, we anticipate that the translation efficiency of the CcdA antitoxin would be at least five-fold higher than the toxin, CcdB (Figure 3D). It is therefore probable that mutations which lead to perturbations in the relative translation efficiencies of the two proteins in the operon context will manifest in an altered phenotype.

### The altered protein ratio does not result from changes in the relative mRNA levels of ccdA and ccdB

To monitor the changes in the mRNA levels, RT-PCR was performed using both CcdA and CcdB specific primers (Figure 3C). A moderate increase in the mRNA levels was observed for most inactive mutants except for positions D71 and L16. Also, ccdA and ccdB mRNA regions were found to be present at similar levels in both WT as well as the mutant samples in this assay. Since the system under study is an operon, it is unlikely that there will be selective changes in the levels of mRNA of one gene relative to the other. It may be important to note that the mRNA quantified is expressed from its native promoter and transcribed by the native cellular *E. coli* RNA polymerase. This is in contrast with other, previous studies that have used T7 RNA polymerase based assays, where the mRNA levels may be mis-estimated as the elongation rate of this polymerase is different from that of *E.coli* RNA polymerase (24). Moderate increase in the ccdAB mRNA levels relative to WT in most inactive mutants is likely a consequence of de-repression of the operon due to decrease in the CcdA: CcdB ratio (19), rather than being the primary cause for differential protein levels (27). This is in contrast to other studies which demonstrate mRNA levels to be directly correlated with protein levels (24). Since these latter studies typically over-expressed the mRNA, phenotypic effects could be biased by limiting *in vivo* tRNA levels.

### Inactive phenotypes are uncorrelated with codon preference

There is no significant over representation of rare codons in both *ccdA* and *ccdB* genes which have an overall GC content of about 51%, similar to the overall GC content of the *E. coli* genome. We analyzed the relative frequency of the mutant codon with respect to the WT codon in the *E. coli* genome, for positions showing synonymous inactive mutations. Inactive phenotypes of all available synonymous mutants correlated neither with codon usage, tRNA abundance nor with a modified codon usage parameter, codon influence (Supplementary Figure 4). This indicates that mechanisms other than codon usage are at play at controlling the synonymous mutant phenotype.

### Assessing folding defects in CcdA mutants through WT CcdB binding probed by yeast surface display

To monitor if the synonymous mutants exhibit rare codon induced folding defects, which may manifest in lower protein yields *in vivo*, we used yeast surface display to monitor the binding of single synonymous CcdA mutants to its partner CcdB. Defects in folding should result in reduced CcdA-CcdB binding. We find that all single synonymous inactive mutants probed, bind with similar affinity as WT CcdA to CcdB (Supplementary Figure 4D). Many mutants in the library show an inactive phenotype in *E. coli* (no growth) but apparently bind well to CcdB as assayed by yeast surface display, and hence are properly folded. Therefore, a decrease in CcdA: CcdB ratio *in vivo*, in *E. coli*, is not due to misfolding of the CcdA mutants.

### Inactive, synonymous CcdA mutants may cause ribosomal pausing

An analysis of bacterial genomes has suggested that SD like sequences within the coding region of genes could be potential ribosomal pause sites and are therefore under negative selection (6) (28). To test if the synonymous mutations in CcdA could generate such potential pause sites, the difference in interaction energies with the anti-SD sequence between single synonymous mutants and the WT sequence with respect to the anti-SD sequence was calculated for a window of 10 nucleotides using the RNAsubopt program in the Vienna RNA package (29). The data (Figure 4) indicates that all inactive mutants had higher (more negative) interaction energy with the aSD sequence than the WT. This additionally suggests that inactive, synonymous CcdA mutants may cause ribosomal pausing. Properties such as changes in GC content upon mutation can lead to such difference in the interaction energies. Also, an increase in base pairing propensities of these nucleotides upon mutation can likely lead to an altered RNA structure, which in turn may cause altered translation efficiencies of the ccdA and ccdB genes in the operon.

**Figure 4:**
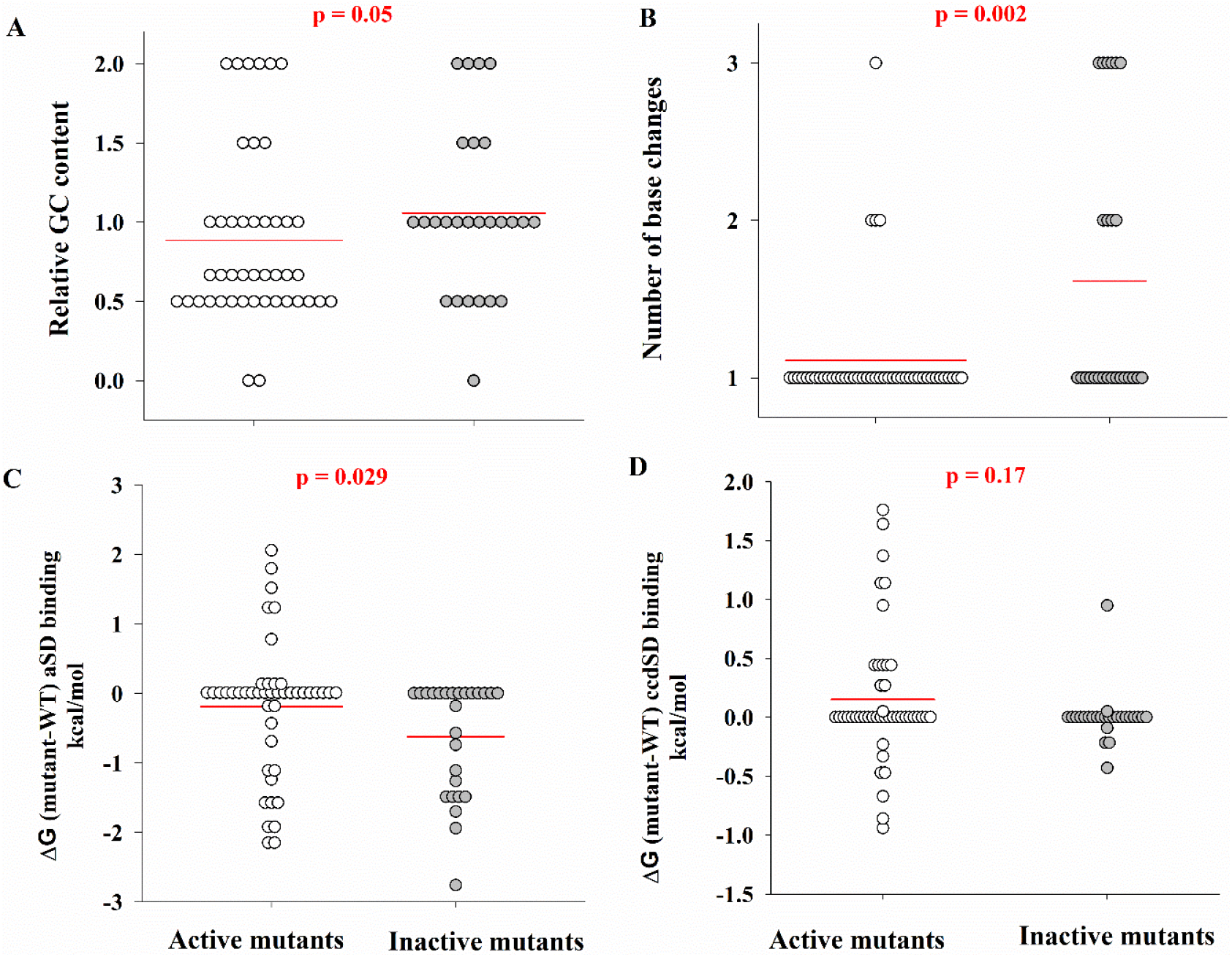
Sequence features of inactive, synonymous CcdA mutants. Single synonymous mutants in CcdA (N=71) were divided into active (N=45) and inactive (N=26) based on a depletion ratio cut-off of 0.3. Mutant distribution is plotted with respect to various sequence based features such as (A) relative GC content, (B) number of base changes, (C) change in binding energy to the anti-Shine Dalgarno sequence on the ribosome (6), and (D) change in the binding energy to the ccdA Shine Dalgarno sequence, relative to the WT, to analyze if the mutation leads to ribosomal pausing or change in accessibility of the ribosome binding site by base pairing with the native SD sequence (30). Statistical significance was assessed by a Mann-Whitney Rank Sum Test. A p-value >0.05 indicates that there is no statistically significant difference between the distributions of the indicated parameter for active and inactive mutants, at 95% confidence interval. The horizontal lines represent the distribution means.

Previously, a Rho-dependent termination mechanism has been proposed for degradation of unprotected mRNAs because of reduced accessibility of the SD sequence upon synonymous mutations (30). This is unlikely for a TA operon, where the expression of downstream toxin gene is essential for an inactive phenotype to manifest. In agreement with this, we did not observe any decrease in the mRNA levels upon synonymous mutations (Figure 3) nor did the mutations lead to significant predicted decrease in SD accessibility (Figure 4).

### Saturation suppressor mutagenesis

We carried out an exhaustive, saturation suppressor screen (31) to obtain additional insights into mechanisms responsible for synonymous inactive mutant phenotypes (Supplementary Figure 5). Analysis of the suppressor mutants demonstrates that synonymous codons at different residues tend to suppress each other (Figure 5). Mutations at residues T8 and V7 suppress more than one single synonymous mutation. Interestingly, the synonymous mutants that suppress other synonymous inactive mutations are individually inactive. This is suggestive of interaction between corresponding nucleotides in the mRNA.

**Figure 5:**
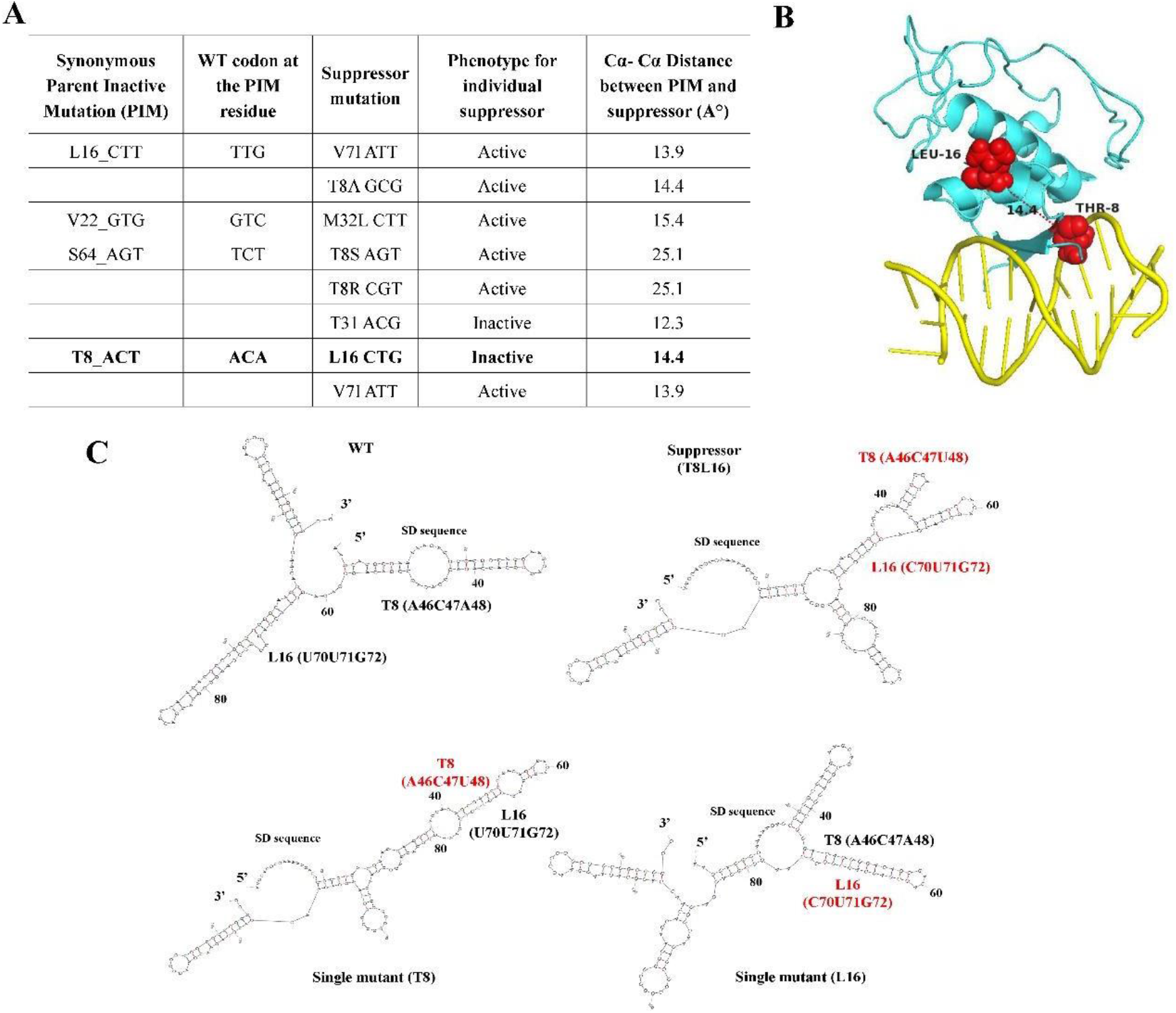
Suppressor analysis for synonymous loss of function mutants. (A) Suppressors isolated for single synonymous inactive mutants in CcdA. (B) T8 and L16 residues mapped onto the CcdA protein structure bound to DNA (PDBid 2H3C). The two residues do not interact with each other and are >14Å apart. (C) Mapping of select synonymous inactive mutations onto predicted mRNA structure. A model of the mRNA structure was generated for the ccdA sequence used for inline probing, for WT, T8_ACU, L16_CUG and T8_L16 suppressor mutant, using the mFOLD server. Bases that show a loss of function phenotype upon synonymous mutation from the sequencing data, and the SD sequence in each of the structures have been boxed. Two of the residues, T8 and L16 in which synonymous suppressor mutations were isolated show unpaired bases in the WT mRNA model structure. In both the T8 and L16 inactive single mutants, the bases encoding T8 mutant (A46, C47, U48) and L16 mutant (C70, U71, G72) are paired to each other, whereas in the suppressor, these bases are paired to other bases in the ccdA sequence forming two distinct stems, making the overall structure different from the individual mutants as well as to the WT.

Consistent with this, two of the suppressors obtained (at residues V7 and T8) for synonymous inactive mutations at L16 residue, show base pairing in the predicted mRNA secondary structures of WT CcdA (Figure 5). The T8 residue is in the DNA binding domain of CcdA and is not proximal to L16 in the DNA-bound protein structure (PDBid 2H3C). This indicates that rescue of WT like mRNA structure by the suppressor in turn results in a WT like CcdA: CcdB ratio *in vivo*. It has been observed that the efficiency of translating a codon is sometimes influenced by the nature of the immediately adjacent flanking codons. A role for triplets of codons in translation has been suggested (32).

We therefore probed the effect of synonymous mutations in the codons adjacent to the inactive mutant codon, however these could not suppress the single synonymous inactive mutation. Of the few transformants obtained in the sensitive strain, many clones had insertions/ deletions making the toxin non-functional, leading to growth under selective conditions. This suggests that altered global RNA structure, rather than adjacent codon effects are responsible for the observed codon specific phenotypes in CcdA.

### Presence of differential secondary structure in the mutant ccd RNAs

We sought to probe the *in vitro* mRNA structure of the mutants and compared it to the WT. Previously, models based on prediction of RNA secondary structure have shown some success in establishing a correlation between alterations in the secondary structure and translation rate differences at the N-terminal end of a gene (12). To experimentally probe possible changes in RNA structure resulting from mutation, *in vitro* transcription was used to generate the first ~150 bases of ccd mRNA which was subjected to alkaline hydrolysis. Base paired regions display protection from hydrolysis (33). In-line probing of WT, suppressor pair, and inactive mutant ccd transcripts all show presence of local secondary structures in the mRNA. Our analysis above suggested that alteration of ccdA SD sequence accessibility was unaltered by downstream synonymous mutations. Consistent with this, we find that the region of the mRNA corresponding to the SD sequence is fully accessible in these different transcripts. Notably, the most prominent differences in RNA structure are seen in a ~36 nucleotide stretch (~G45 to G81) which encompasses the T8 and L16 codons. These are paired to each other in models of two individual single mutants, T8_ACU and L16_CUG (Figure 6). While transcripts for the individual T8 and L16 mutations look distinctly different in structure from that of the WT, the suppressor double mutant appears to restore the local secondary structure to more closely resemble the WT. The L16 inline probing data also clearly reveal a stem region comprising of segment G45 to G81 which is not visible in other mutants. Although there is no clear correlation between the protection seen in the inline probing with base paired regions in the predicted structures, both approaches collectively suggest that synonymous mutations likely cause a change in mRNA structure leading to a change in CcdA: CcdB ratio, resulting in cell death.

**Figure 6:**
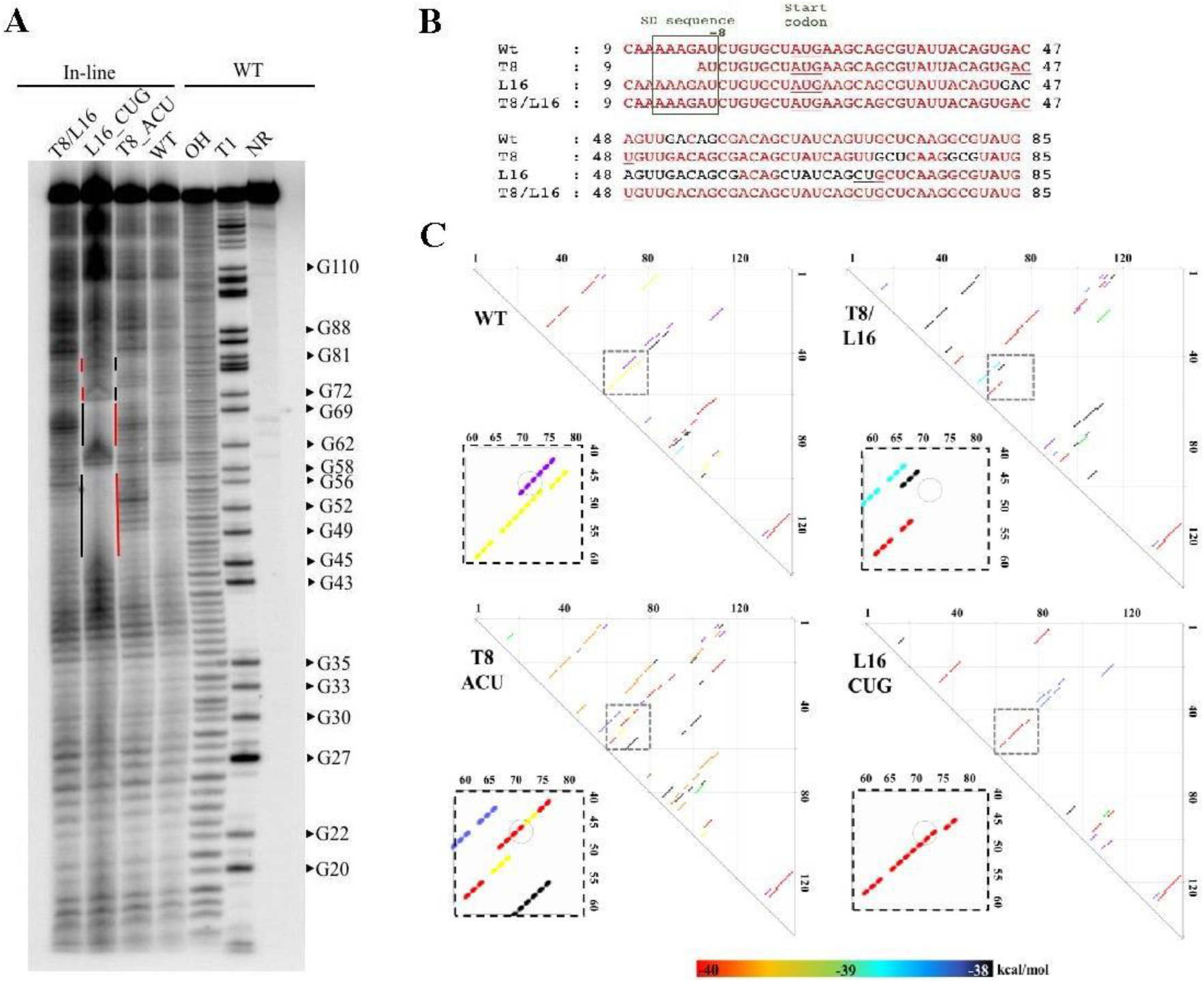
Probing of WT and mutant ccd RNA structure. (A) In-line probing of the ccd mRNAs. Vertical lines to the left of a lane indicate regions of structural differences, black for protected (structured) regions, red for unprotected (flexible) regions. Protection patterns for the WT and suppressor (T8/L16) double mutant are more similar to each other than to the patterns observed for the two inactive single mutants, T8_ACU (nucleotide positions 46,47,48) and L16_CUG (nucleotide positions 70,71,72). NR is the no reaction negative control. Base positions were annotated from the G specific RNAse T1 cleavage and the OH ladders. (B) Nucleotide sequence alignment of the annotated ccdRNAs from the in-line probing analysis showing the changes in protection with respect to the WT ccd RNA sequence. The bases in red are unstructured and not protected and the ones in black are structured and protected in the probing reaction. (C) Energy dot plots for WT, T8_ACU, L16 CUG and T8_L16 mutants taken from the Mfold server. The upper triangles depict the base pairs within the sequence. The energies of pairing have been color-coded. The region encompassing the bases encoding T8 mutant (A46, C47, U48) and L16 mutant (C70, U71, G72) has been boxed. Pairs between the bases A46, C47, U48 and C70, U71, G72 bases is encircled.

### Effect of designed synonymous mutations in another *E. coli* TA operon, RelBE

To test if insights from the ccd operon had predictive value, the RelBE system was used as a test case. Unlike the plasmidic ccdAB system, relBE is a chromosomal toxin-antitoxin system expressed as a part of the relBEF operon (34). The RelE toxin is a site- specific, ribosome dependent mRNA endonuclease that leads to bacterial growth inhibition. Mild overexpression of RelE also leads to increase in the persister cell frequency (26). While unrelated in sequence, the RelB protein from *E. coli* is like the CcdA antitoxin in both fold and thermodynamic properties and inhibits the action of RelE by forming a tight complex. The RelBE complex acts as a transcriptional repressor and auto- regulates its own expression (34). Homologs of this system are found in many bacterial species including pathogens like *Haemophilus influenzae*. An analysis of the ribosome profiling data for the *E. coli* K12 MG1655 strain indicate that the translation efficiency for the relB gene is indeed 3.6-fold higher than the relE gene, although both the codon and tRNA adaptation indices are similar for both genes, similar to what we inferred in the CcdAB system. From the ccd study, we found that most synonymous inactive mutants were three base substitutions. We attempted to elicit an inactive phenotype upon introduction of synonymous mutations at single codons in the RelBE operon. We designed and tested seven single synonymous substitutions in relB, including all possible three base substitutions in the serine codons, three representative two base substitutions and one control mutation at the 77^th^ residue of relB that strengthens the SD sequence of RelE, thereby leading to an increase in RelE levels and inhibition of cell growth (Figure 7A). These mutations were to both optimal and rarer codons. Using a similar screen of cell death vs. growth as in the case of CcdAB system, we monitored the growth of single synonymous mutants in the WT (*E. coli* BW25113 *relBE*) as well as in the toxin deleted (*E. coli* BW25113 ΔrelE) strain. Most mutants had a decreased growth in the WT strain, indicating that the designed synonymous mutations indeed result in a loss of function phenotype (Figure 7B). This also indicates the existence of a relatively general phenomenon wherein a change in RNA sequence upon synonymous mutations can have phenotypic effects that are unrelated to the change in codon usage frequency upon mutation, at least in toxin-antitoxin type operon systems, where small perturbations in the system can compromise cellular fitness. Furthermore, these studies highlight the utility of synonymous mutations in perturbing *in vivo* expression levels without altering protein sequence, to result in desired phenotypes.

**Figure 7:**
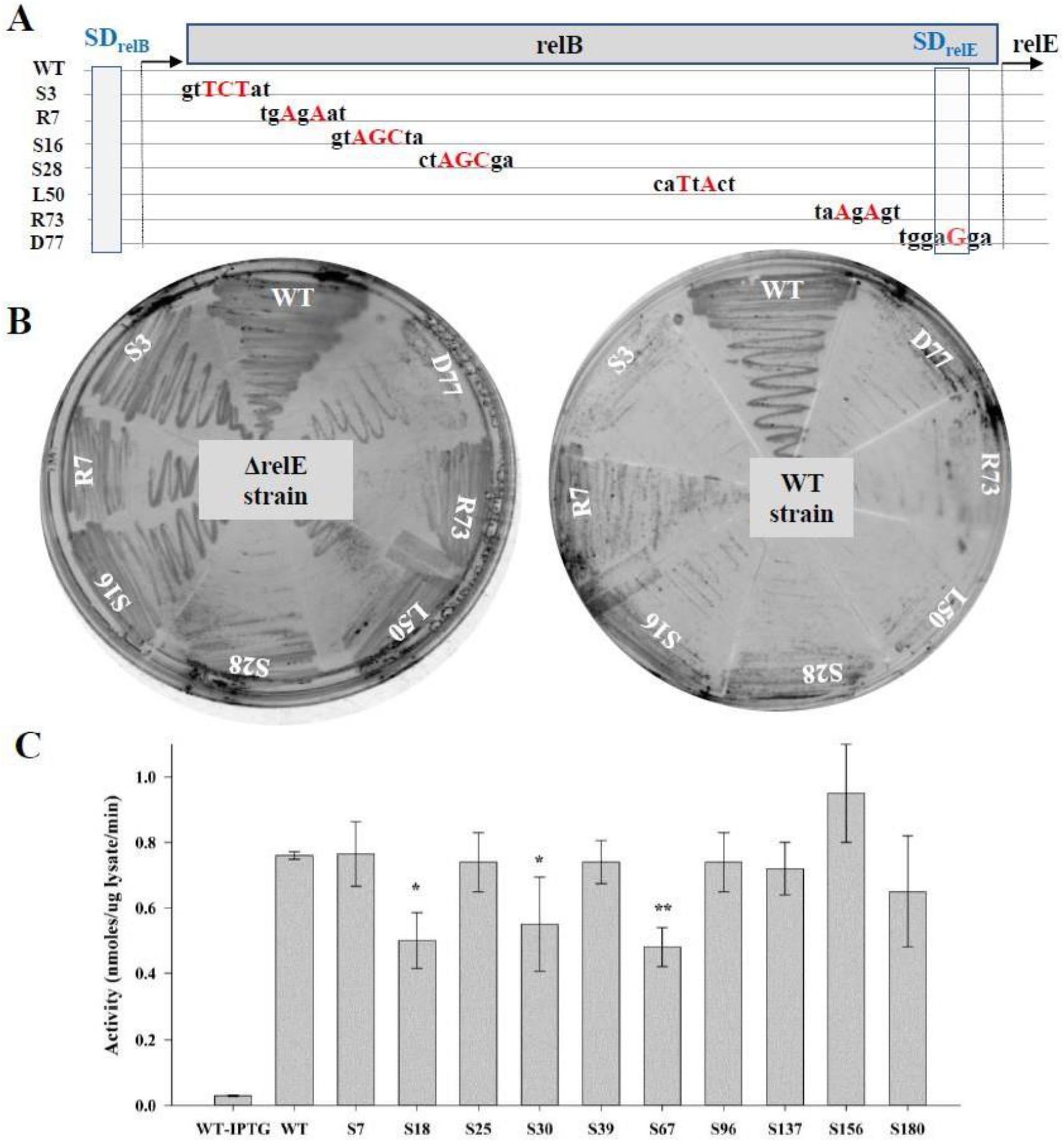
Phenotypic effect of designed single synonymous substitutions in RelBE operon and LacZ genes. (A) Location and identity of various synonymous mutations made in the relB gene in the operon context. Mutant bases are highlighted in red. Regions encompassing the SD sequence for both relB and relE genes have been marked. A single base substitution at the D77 residue was used to increase the strength of the SD sequence for relE and was used as a control mutant. (B) Growth of single synonymous mutants in RelB overexpression (*E. coli* BW25113 ΔrelE) and WT (*E. coli* BW25113 *relBE*) strains. Following transformation in the indicated strains, single colonies were streaked into sectored plates. Most mutants showed a decreased growth in the WT strain indicating that the synonymous mutations in the relB gene results in a loss of function phenotype. (C) Activity of single synonymous mutants in the α-peptide encoding region of LacZ. Error bars indicate standard error. ‘*’ indicate significant differences from the WT (* p-value <0.05 and ** p value <0.01, paired t-test).

### Synonymous mutations in a non-operonic context: Test of generality using β-galactosidase enzyme as a model

We next tested the effect of three-base synonymous substitutions in a non-TA gene model. We selected the α-peptide of the β- galactosidase enzyme as a facile activity assay exists and there is extensive prior mutational data for the β- galactosidase gene, including mutations that alter activity by affecting mRNA structure (35, 36). Ten single mutants at serine codons, distributed throughout the gene, were made. Mutants were transformed in *E.coli* XL-1 Blue strain and activity was monitored using a colorimetric assay. Three out of ten mutants have significantly lower activity than WT (Figure 7C). As before, we did not find any correlation between codon usage and mutant growth. Most earlier studies on β- galactosidase gene have found synonymous mutations to be near neutral, with a majority of mutants being one or two base substitutions (35, 36).

## Discussion

Our studies on the *ccdA* gene in an operon context using saturation mutagenesis coupled to deep sequencing revealed that several synonymous substitutions along the length of ccdA can lead to a loss of function phenotype. The present work demonstrates that there can be strong selection for specific synonymous codons, complicating interpretation and use of dN/dS ratios. The frequency of synonymous mutations that affect phenotype in our system is large, with ~38% (28 of 73) of all assayed synonymous mutations having a loss of function phenotype. Many non-synonymous mutations also show codon specific phenotypes. Quantitative proteomics results indicate that for loss of function mutants, the *in vivo* CcdA: CcdB ratio is lower than WT leading to cell death. The data suggest that synonymous mutations in CcdA cause a change in mRNA structure and thereby leading to a change in the CcdA: CcdB ratio and affect operon auto-regulation. The ccd operon used in the present experiments differs slightly in sequence from the one on F-plasmid (Figure S1) resulting a slightly reduced CcdA:CcdB ratio (Figure 3B), thus enhancing sensitivity to mutations. This ccdAB system allows reporter independent monitoring of *in vivo* phenotypes, that are highly sensitive to small perturbations in the regulation of the system. The insights obtained were used to design synonymous loss of function mutations in the unrelated relBE operon and in the gene encoding the lacZ α-peptide. Earlier studies which observe a strong influence of codon rarity and codon pair bias on protein level (24, 37), typically used over- expressed proteins in a non-physiological context. In natural contexts, many bacterial genes are organized in operons to ensure co-regulation of genes involved in similar pathways and functions. In the special case of toxin-antitoxin operons, fine tuning of the relative amounts of antitoxin and toxin is essential and directly linked to bacterial fitness. Given the abundance and diversity of these TA systems in many bacterial species, including the pathogen *Mycobacterium tuberculosis*, it is reasonable to think that the cell must have evolved multiple mechanisms to ensure their stringent spatial and temporal regulation. Analysis of whole genome RNA-seq and ribosome profiling data available for the *E.coli* genome indeed indicate that while the mRNA levels for most type II TA systems were similar for the toxin and the antitoxin, the antitoxin protein levels can be upto two fold higher than the toxin under normal growth conditions (16). We find that these relative protein levels can be perturbed through changes in mRNA structure due to synonymous mutations. A few recent reports have highlighted widespread selection of local mRNA structure in the coding regions of viral RNAs, predominantly through computational analysis of relative folding free energies of the different RNA regions (38) and identifying sites of suppression of synonymous codon usage from a multiple sequence alignment (39). A recent finding suggested that the order of the genes in operons is reflective of the assembly of the gene products (40). Given the conservation of the proximity of the toxin binding domain on the antitoxin, to the toxin coding region, it is reasonable to think that the folding and binding of the antitoxin and toxin may be co-translational *in vivo*. This may pose further constraints on the selection of codon sequence that will be probed in future studies. While it was earlier thought that most mRNAs are relatively unstructured, recent genome-wide studies have shown varying degrees of structure in *E. coli* mRNAs (13). However, the functional implications of these findings, and the sensitivity of these structures to point mutations have not been studied. The current study confirms that these structures are indeed sensitive to mutations and that synonymous mutations can have large phenotypic effects, in a natural context. Mutational tolerance inferred from laboratory-based mutational studies is typically much higher than observed natural sequence variation. However, the present study suggests that this may be a consequence of the artificial selection conditions, overexpression and other factors. Thus, genes are likely more sensitive to mutation than inferred from previous saturation mutagenesis studies (10, 41).

## Acknowledgments

We acknowledge funding for infrastructural support from the following programs of the Government of India: DST FIST, UGC Centre for Advanced study, Ministry of Human Resource Development (MHRD), and the DBT IISc Partnership Program. The funders had no role in study design, data collection and interpretation, or the decision to submit the work for publication. This work was funded in part by a grant to RV from the Department of Biotechnology, grant number-BT/COE/34/SP15219/2015, DT.20/11/2015), Government of India and grants from DST and MHRD.

## Author Contributions

R.V. and K.G. designed the experiments, except for the quantitative proteomics (S.V.M. and H.G.), RNA inline probing (J.P. and A.R.) and yeast surface display studies (S.C.). A.A helped in acquiring FACS data. S.K. processed the Illumina data. P.K performed the experiments with LacZ. R.V. and K.G. analyzed the overall data and wrote most of the manuscript with critical inputs and review from all other authors.

## Methods

### Preparation of a single-site saturation mutagenesis library of CcdA

Mutagenic primers for all 71 positions (2nd to 72nd) of CcdA were designed such that the degenerate codon (NNK) was at the 5’ end of each 21 bp forward primer. A non- overlapping adjacent 21 bp reverse primer, along with the forward primer, was used to amplify the entire pUCccd vector by inverse PCR methodology (22). The primers were obtained in 96-well format from the PAN Oligo facility at Stanford University. A master-mix was made for doing the PCR for all positions in a 96 well format as per the manufacturer’s instructions (Phusion DNA Polymerase). Following quantification, an equal amount of PCR product (~200ng) at each position was added to get a pool of PCR products of all required fragments. Gel-band purification of the pooled PCR product at the required size (~3.6 Kb) was done using a Fermentas GeneJET™ Gel Extraction Kit. After purification, pooled PCR product was phosphorylated, followed by ligation. The ligated product was transformed into high efficiency (10^9^ CFU/μg of pUC57 plasmid DNA) electro-competent *E. coli* Top10Gyr cells. 5μg of the total ligated PCR product was transformed into 2mL of competent cells (4 aliquots of 500μl each) and the cells were plated on LB agar plates containing 100μg/mL ampicillin, for selection of transformants. Plates were incubated for 18-20 hrs at 37°C. Around 2000-3000 colonies obtained on each plate were washed off from a total of 8 LB agar plates (150mm diameter) into a total volume of 100 mL of LB media. Pooled plasmid was purified using a Qiagen plasmid maxiprep kit.

### Sample preparation for deep sequencing and data processing

Pooled, purified plasmid samples from each condition were PCR amplified with primers containing a six-base long Multiplex Identifier (MID) tag. 270 bp long PCR products, containing the full ccdA gene, were pooled, gel-band purified and sequenced using Illumina Sequencing, on the MiSeq platform. Sequencing was done at Macrogen, Korea. The sequencing was done for two biological replicates. The initial quality of the sequencing data was assessed using FASTQC software. Further analysis was performed using in-house Perl scripts. Sequencing data was analyzed by assigning each read to a particular “bin” based on its MID tag. The downstream primer sequence was used to identify forward and reverse reads in each bin. Only those forward and reverse read pairs which overlap with each other, and together cover the entire *ccdA* gene length, were considered for analysis. The reads were aligned with the wild-type sequence using the Water program of the JEMBOSS package (42). Reads with insertions, deletions and multiple mutations were omitted, and only mapped single-mutants were used for further data analyses.

### Assignment of mutant phenotypes based on the deep sequencing data

Read numbers for all mutants at all 71 positions (2-72) in CcdA were analyzed. Mutants having <5 reads in the resistant strain were not considered for analysis. The total number of reads in different conditions was calculated. Read numbers for each mutant at a given condition were normalized to the total number of reads in that condition. Depletion ratio was defined as the ratio of the normalized reads in the sensitive strain to the normalized reads in the resistant strain, for a given mutant. For each residue in the CcdA protein, reads for all the 32 possible mutant codons were analyzed in the sensitive vs. resistant strain and phenotypes were assigned. Positions showing codon specific mutation effects were analyzed separately.

### Construction of double mutant library to identify suppressors for synonymous inactive mutants

#### Saturation suppressor libraries

Single CcdA synonymous inactive mutants were introduced into the CcdA library using the inverse PCR approach described above and transformed first in the resistant strain, *E. coli* Top10Gyr, to get the entire double mutant library. Pooled plasmid from this library was isolated and retransformed in the sensitive strain, *E. coli* Top10 to select for suppressor mutants. 96 clones were streaked on a plate and sent for Sanger sequencing at Macrogen to determine the identity of the suppressors.

#### Codon pair libraries

Double mutant libraries were constructed in the background of 6 selected single-site inactive, synonymous mutants to introduce all synonymous mutations both in the upstream and downstream codon, using the inverse PCR strategy, described above. Pooled mutant plasmids for each position were extracted from the resistant strain and transformed into the sensitive strain to obtain suppressors.

### *In vivo* activity of individual single-site CcdA mutants

Selected single-site synonymous mutants of CcdA were constructed and sequence confirmed. These plasmids were retransformed into the sensitive and the resistant strain and plated on LB agar plates and grown at 37°C for 16 hrs. CFU’s for the transformants obtained in the two conditions were counted to confirm their *in vivo* activity.

### Growth kinetics upon over-expression of strains transformed with single synonymous mutants

The CcdAB coding sequence was synthesized and cloned in the pBAD24 expression vector at GeneArt (Germany) to drive the expression of CcdA and CcdB from the pBAD promoter, which is inducible by arabinose. A set of 11 single synonymous mutants of CcdA were made in the above construct. 200 μl of mid-log cells (OD_600_ of 0.4) were grown in a microtitre plate. Growth was monitored for 5 hours, by conventional OD_600_ measurements.

### Monitoring the relative levels of CcdA and CcdB using a quantitative proteomics approach

#### Selection of proteolytic peptides from CcdA and CcdB for MRM experiments

In order to identify proteolytic peptides, *E. coli* lysates were resolved on SDS-PAGE and gel bands corresponding to CcdA and CcdB were excised and subjected to proteolytic digestion using trypsin. Peptide digests were analyzed on Thermo Orbitrap Fusion^TM^ Tribrid^TM^ Mass Spectrometer interfaced with Easy-nLC 1000 nanoflow liquid chromatography system (Thermo Scientific, Bremen, Germany). Peptides were loaded onto a trap column (75 μm × 2 cm, Magic C18AQ, 5 μm, 100 Å, Michrom Biosciences Inc., Auburn, CA) using 0.1% formic acid at a flow rate of 3 μl/min. The peptides were then resolved on an analytical column (75 μm × 20 cm, Magic C18AQ, 3 μm, 100 Å, Michrom Biosciences Inc, Auburn, CA) at a flow rate of 350 nl/min using a linear gradient of 5-35% B (0.1% formic acid in 95% acetonitrile) over 18 min and a run time of 30 min. The MS and MS/MS scans were acquired at a mass resolution of 120,000 and 30,000 at 200 m/z, respectively. Full MS scans were acquired in the m/z range of 350-1550. Precursor ions with single charge or unassigned charge were rejected. Dynamic exclusion of fragmented precursor ions was set to 30 sec. Precursor ions were isolated using a quadrupole mass filter with isolation width of 2 m/z. Higher energy collision dissociation was used as the fragmentation method with 32% normalized collision energy. Proteome discoverer (version 1.4) software suit (Thermo Fisher Scientific, Bremen Germany) was used to analyze the data. 3 peptides were selected for CcdA and 2 for CcdB. Peptide ITVTVDSDSYQLLK and LFVDVQSDIIDTPGR was used for quantitation of protein CcdA and CcdB, respectively while others were used for identification purposes only.

#### Mass spectrometry protein sample preparation

To quantify the levels of CcdA and CcdB proteins in the *E.coli* lysate, 10 ml culture of *E. coli* Top10Gyr cells transformed individually with different CcdA mutants were grown till saturation (OD_600_ of 1.0) under shaking conditions at 37°C, 180 rpm. Cells were pelleted, washed twice with PBS, and lysed by sonicating in lysis buffer (2% SDS in 100mM triethyl ammonium bicarbonate (TEABC) buffer, pH 8.5). The lysates were clarified by centrifugation at 12,000 g for 30 min at 4°C. Proteins were precipitated with ice-cold acetone to remove SDS and re-constituted in 100mM TEABC. Bicinchoninic acid (BCA) assay kit (Thermo-Scientific) was used to estimate protein concentration. An equal amount of protein from each sample was reduced with 10mM DTT at 60°C for half an hour. The samples were cooled to room temperature and proteins were alkylated by adding iodoacetamide at a final concentration of 20mM and incubating at room temperature in the dark for 15 min. Proteins were subjected to proteolytic digestion overnight by adding sequencing grade trypsin (Promega) at an enzyme:substrate ratio of 1:20. For LC-MS-MRM experiments, samples were spiked with stable isotope labeled peptides (SpikeTides™ TQL peptides were purchased from JPT Peptide Technologies, Germany) before adding trypsin. After overnight incubation, the reaction was quenched with 10% formic acid and centrifuged at 12,000 g to remove any precipitate. Sample clean-up was performed using reverse phase spin columns (Harvard apparatus) according to the manufacturer’s protocol.

#### SRM/MRM analysis

LC-MS/MRM analysis was performed with an electrospray ionization-triple quadrupole mass spectrometer (QTRAP 6500; SCIEX) coupled with HPLC system (Agilent 1260 systems; Agilent Technologies, Santa Clara, CA). The peptides were resolved on a reverse phase C18 column (Agilent Poroshell 120 EC-C18 2.7μm, 4.6×50mm) at a flow rate of 350 μl/min and a run time of 10 min. The LC method used was as follows: 10% - 30% solvent B from 0-3 min, 30%-50% solvent B from 3-5 min, 50%-95% solvent B from 5-7 min and 95%-5% solvent B from 7-10 min. The data was acquired in SRM/MRM mode using a dwell time of 25 msec. Each target peptide and corresponding SIL peptide were measured. Standard curves were generated by spiking different amounts of SIL peptides (0.5, 0.1, 0.25. 0.5, 1, 5, 10, 20 pmol) into a constant amount of *E. coli* digest background.

#### LC-MS/MRM data analysis

MRM data was analysed using Skyline 3.5 (43) and composite peak AUCs were calculated after Savitsky-Golay smoothening. Each transition was inspected and XICs for each transition was generated. Response curves were generated by plotting the ratio of heavy:light peptide over different concentrations of spiked heavy peptide. Linear regression analysis was carried out to determine the linearity of the response curve. Also, to validate the analytical method, regression analysis of variance (ANOVA) of the linear regression data measurements (at a 95% confidence level) was performed. In addition, to illustrate the reliability of LC-MS/MRM analysis, the %RSD was calculated which was within 20% in all cases. Peaks corresponding to endogenous target peptides were identified based on the retention time of the corresponding SIL peptide as it co- elutes with the endogenous peptide. Welch’s test was used to determine the statistical significance of abundance differences between the wild type and mutants.

### Estimation of ccd specific mRNA levels using qRT-PCR

For RNA isolation and quantitation, 3 ml of culture was grown for each of the mutants of ccdA transformed in *E.coli* Top10Gyr. Cells were grown to saturation under shaking conditions at 37°C, 180rpm, pelleted and RNA was extracted by the RNAsnap method (44). Chromosomal and plasmidic DNA were removed by treatment with 2 units of DNase1 for 2 hours at 37°C, followed by ethanol precipitation of RNA. RNA was quantified by nanodrop spectrophotometric estimation, and quality was assessed by agarose gel electrophoresis prior to downstream processing. 2μg of total RNA was taken in a sterile, RNase-free microcentrifuge tube and 0.5μg of random hexamers was added to serve as primer for cDNA synthesis, in a total volume of 15μl in water. The mixture was heated to 70°C for 5 minutes to melt secondary structure within the template. The tube was cooled immediately on ice to prevent secondary structure from reforming. To this mix, 200 units of Moloney Murine Leukemia Virus (M-MLV) Reverse Transcriptase was added, along with dNTPs, M-MLV 5X Reaction Buffer and 25 units of Ribonuclease Inhibitor. The reaction mix was incubated for 60 minutes at 37°C. cDNA was directly used for quantitative PCR (Q- PCR) analysis. Q-PCR was set up with ccd gene specific primers using the cDNA template in a Bio-Rad iQ5 machine. A reaction with no reverse transcriptase was used as a negative control. 16S rRNA was used as the internal positive control for total RNA.

### Experimental determination of transcription start site

Total RNA was extracted from *E.coli* Top10GyrA cells transformed with pUCccdAB plasmid using RNA SNAP method (44). First strand cDNA synthesis was done using gene specific reverse primer (R1) phosphorylated at the 5’ end. The cDNA was circularized using T4 DNA ligase. Circularization brings the 5’ end of the transcript (TSS) next to the 5’ end of R1 primer. Using this as a template, PCR amplification using internal forward and reverse primers (IF1 and IR1) was performed to generate a fragment containing the TSS and R1 sequence. The fragment was sanger sequenced. Base preceding the 5’ end of the R1 primer was assigned as the transcription start site for ccdAB transcript.

### In-line probing of the Ccd mRNA

#### Preparation of RNAs

The 167 bp region of the ccdAB operon from the experimentally determined transcription start site till amino acid 41 of the ccdA gene was used for the study. ccd DNA template containing a flanking T7 promoter (incorporated by using forward primer at the 5’ end) was amplified using PCR and used to generate RNA by *in vitro* transcription. 25μl of *in vitro* transcription reactions contained ~2.0 μg of DNA template 80 mM HEPES, pH 7.5 at 25°C, 40 mM dithiothreitol (DTT), 24 mM MgCl_2_, 2 mM spermidine, with 2.5mM of each nucleoside triphosphate (NTP) and recombinant T7 Polymerase. RNA was purified using 6% denaturing (8M Urea) PAGE. The 150-nucleotide long RNA was excised from the PAGE gel. The gel was minced, and RNA was recovered passively by incubating gel pieces at room temperature for 2 hours in crush soak buffer (200mM NaCl, 10mM Tris- HCl, pH=7.5, 1mM (Na)_2_EDTA, pH=8.0). The eluted RNA was ethanol precipitated and the pellet was re- dissolved in nuclease free water.

#### Labeling of RNAs with [γ-^32^P] ATP

Purified RNAs were treated with alkaline phosphatase (Calf Intestinal Phosphate from New England Biolabs) at 50°C for 15min followed by heat inactivation at 95°C for 2.5min, to remove the 5’ phosphate. After phenol: chloroform: isoamyl alcohol (25:24:1) extraction, the RNA was ethanol precipitated in the presence of 20μg of Glycogen (concentration- 20mg/mL, ThermoScientific). The RNA pellet was dissolved in nuclease free water. About 20picomoles of CIP treated RNA was 5’ end labeled with 50μCi of [γ-^32^P] ATP (10mCi/ mL, BRIT, India) by incubating with T4 polynucleotide kinase (New England Biolabs) at 37°C for 35 min. The labeled RNA was purified using 6% denaturing (8M Urea) PAGE by crush soak as described above.

#### In-line Probing reaction

The 5’ γ-^32^P labeled RNAs (about 400 CPS) were incubated in In-line buffer (50mM Tris- HCl pH=8.3, 100mM KCl, 20mM MgCl_2_) at room temperature for 44 hrs. Non-reacted RNA along with T1 and OH controls were generated from the RNAs to annotate the nucleotide positions. The T1 cleavage ladder was generated by digesting 400 CPS of respective RNAs at 50°C for 20 min with RNaseT1, (1U/ μL) in 25mM Sodium Citrate buffer pH 5.0 and Urea loading buffer (540mM Tris, 54mM Borate, 6mM EDTA (pH=8.0), 10.8M Urea, 12% Sucrose, 0.06% SDS as final concentration). The reaction was quenched by adding Urea loading buffer (270mM Tris, 27mM Borate, 3mM EDTA (pH=8.0), 5.4M Urea, 6% Sucrose, 0.03% SDS, Xylene Cyanol and Bromophenol blue). The OH cleavage ladder was generated by incubating 400 CPS of the respective RNAs in 50 mM Na2CO3 pH 9.0, 1 mM EDTA, at 95°C for 5 min, followed by addition of Urea loading buffer. Reactions were resolved on a 10% denaturing (8M Urea) PAGE gel run at 40 watts for ~3hrs. Following electrophoresis, the gel was dried on a gel drier at 80°C and the gel post drying was exposed to a screen and developed using a Phosphorimager. Base positions were assigned from cleavage ladders and structured and unstructured regions were annotated.

### Computational prediction of mRNA secondary structure

The initial 150 bp of the *ccdAB* transcript were submitted to the mFOLD (45) for prediction of secondary structure. All energy parameters were set at default values. Detailed output was obtained in the form of structure plots with reliability information, single strand frequency plots and energy dot plots. Local mRNA secondary structural elements encompassing the region of synonymous inactive mutations were further analyzed.

### Calculation of interaction energies for synonymous mutants in ccd mRNA

The difference in the interaction energy between single synonymous mutants in CcdA and the WT sequence, with the consensus anti-Shine Dalgarno (aSD) sequence (5’ CCUCCUAU 3’), was calculated for a window of ten nucleotides using the RNAsubopt program from the RNAVienna package 2.4.3 (29). The average energy difference for each mutant across these ten windows was compared for all the available synonymous active and inactive mutants. Such aSD like sequences are known ribosomal pause sites (6). Similarly, the difference in the interaction energy between single synonymous mutants in CcdA and the WT sequence with the ccdA SD sequence (5’ AAAAAGAU 3’) was also calculated.

### Yeast surface display of CcdA library to probe CcdB binding

The CcdA libraries, individually recovered both from the resistant and the sensitive strains, were amplified from the pUCccd vector and recombined into the yeast surface display vector, pETcon (Addgene plasmid # 41522) between NdeI and XhoI sites, using yeast homologous recombination in *S. cerevisiae* EBY100 strain. Positive clones, sufficient in number to cover the library diversity, were obtained on selective SDCA plates. The transformant pool obtained were grown in liquid SDCAA media for 48 hrs and stored in aliquots of 10^9^ cells per ml in SDCA media containing 20% glycerol at −70°C. The same methodology was used for cloning WT and individual synonymous mutants of CcdA into pETcon. The CcdA coding region was cloned as a C-terminal fusion to the yeast Aga2p protein with a C-terminal c-myc tag under control of the GAL promoter. The library was displayed on the yeast surface, as described (Chao et al, 2006). Cells were induced and grown in SGCA media containing 2% galactose at 30°C for 16 hrs. The surface expression of CcdA was monitored by incubation of 10^6^ cells with anti-c-myc antibody raised in chicken (1:400 dilution), that binds to the Myc tag at the C-terminus of CcdA. Anti-chicken IgG conjugated AlexaFluor-488 (1:300) was used as the secondary antibody. CcdB binding to the surface expressed CcdA was assessed by incubating cells with a fixed concentration of 2nM biotinylated CcdB (or varying concentrations between 0.1pM and 200nM for titration experiments) and binding of streptavidin conjugated AlexaFluor-633 secondary antibody (1:2000) to biotinylated CcdB was monitored. Labelled cells were analyzed on a BD FACSARIA III. Double plots for mean fluorescence intensities for both expression and binding for the CcdA library as well as for single synonymous mutants were analyzed to monitor differences with respect to the WT.

### Monitoring the effect of single synonymous mutations in the RelBE operon

#### Design of mutants

The WT sequence for the RelBE operon from *E.coli* K12 MG1655 strain was retrieved from the NCBI database. The entire transcription unit of 910 bp (46) including another gene relF, which is expressed as a part of the same operon, was cloned in the EcoRV site of the pUC57 vector at Genscript. Seven different constructs having synonymous mutations in the relB gene were synthesized with a stop codon in the relE toxin gene to ensure normal cell growth even after inactivation of the antitoxin. Based on our results on the CcdAB system, either two or three base substitutions were made in order to have an inactive phenotype. Three base synonymous substitutions are only possible with serine codons. Three base synonymous substitutions were made in three of the serine codons, S3, S16 and S28, all at the N-terminal half of the relB coding sequence. Three representative two base substitutions at positions R7, L50 and R73 were also tested. A single base substitution at the D77 residue was made to increase the strength of the SD sequence for relE and was used as a positive control mutant. The introduced stop codon in the relE gene was reverted using the inverse PCR strategy described earlier (22)

#### Phenotype testing

Due to absence of a strain that is resistant to toxin action, we employed a toxin deletion strain (*E.coli* BW25113 ΔrelE) for the propagation of mutants. By virtue of this deletion, the WT chromosomal operon is de-repressed, leading to an excess of antitoxin in the cell, relative to the WT strain (*E.coli* BW25113 relBE). The inverse PCR products for individual mutants were phosphorylated and ligated, as described previously. Ligation mixes were individually transformed into the *E. coli* BW25113 ΔrelE strain, to recover the plasmid. Single clones were isolated, and sequence confirmed. Equal amounts of the mutant plasmids were transformed in *E. coli* BW25113 WT strain (having chromosomal relBE genes intact) to test their phenotype as well as in the *E. coli* BW25113ΔrelE strain as a control. These were then streaked onto LBamp agar plates and grown for 12 hours at 37°C to monitor differences in growth.

### Monitoring the effect of single synonymous mutations in the β- galactosidase gene

The sequence of the α- peptide from β- galactosidase gene was taken from pBluescript II KS+ vector. Ten mutants, each having three base substitutions at serine codons were made at Genscript. The mutants were transformed in *E. coli* XL-1Blue strain. Transformants were selected on LBamp (100μg/ml) plates. Single colonies were picked and grown overnight in LB media. 2 ml culture was pelleted down and resuspended in PBS. Resuspended cells were lysed by sonication, followed by centrifugation at 12000 rpm. Supernatant was transferred to a fresh tube. 20μl of cell lysate was added to a 96 well microtiter dish. To the cell lysate, 120 μl of buffer containing 100mM NaH_2_PO_4_, 10mM KCl, 1mM MgSO_4_ and 50mM β-Mercaptoethanol was added. After 10 minutes of incubation at 37°C, 50 μl of ONPG substrate (4mg/ml) was added. Plate was covered and incubated at 37°C, until the solution turned yellow. Reaction was terminated by adding 90 μl of 1M Na_2_CO_3_ and microtiter plate was read at 420nm. β-Galactosidase activity was expressed as nmoles of β-galactose formed per minute per mg of lysate at 37°C (47).

