## Supplementary data for "Loss of function from widely distributed, synonymous mutations at single codons"

### Supplementary figures

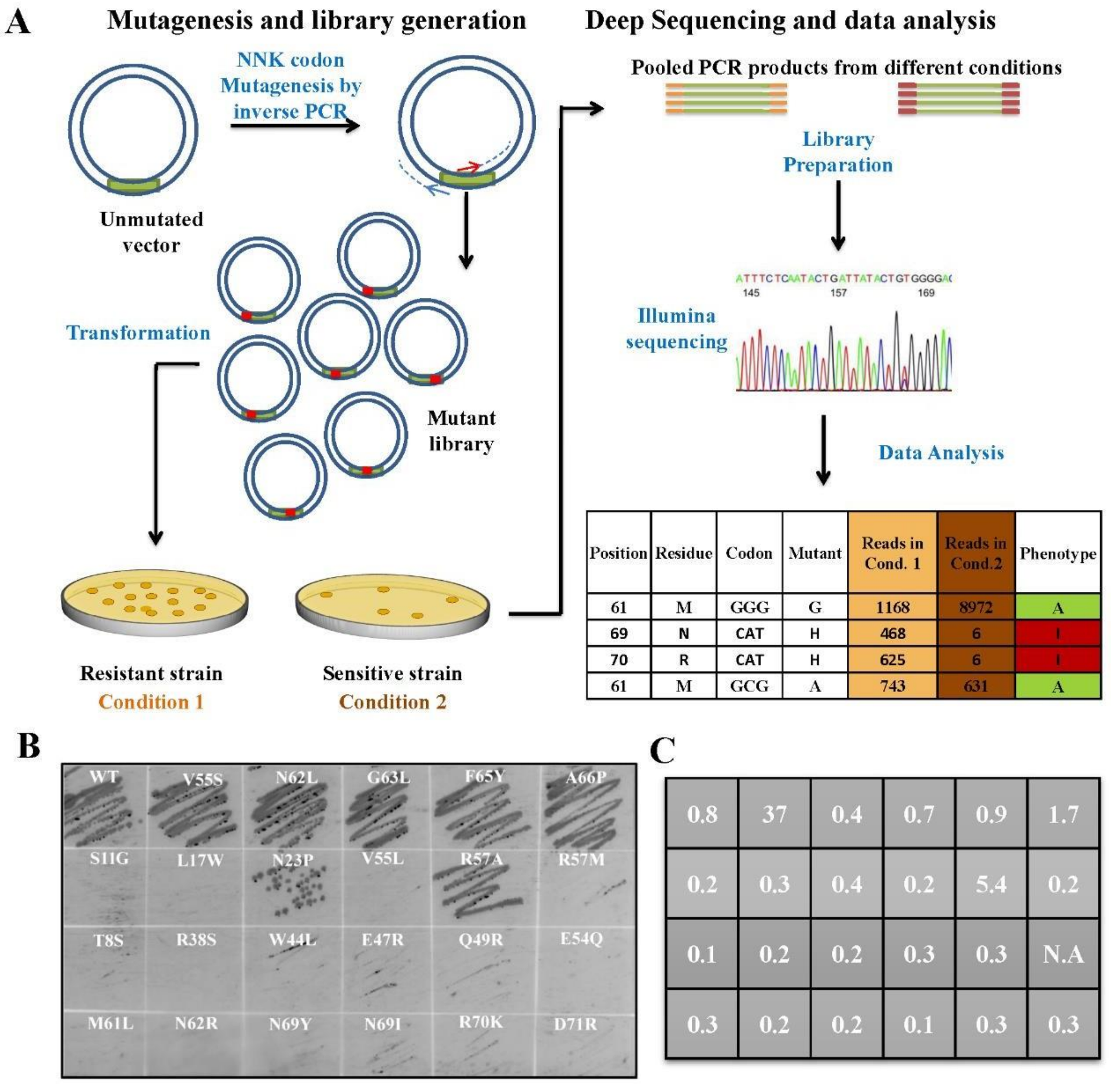


**Figure S1: Saturation mutagenesis coupled with deep sequencing to infer mutational sensitivities.** (A) Single site- saturation mutagenesis libraries are generated using inverse PCR based methodology that involves use of non-overlapping primers with the mutant NNK codon at the 5’ end of the forward primer. The mutants in libraries are assayed by transforming into resistant (condition 1) and sensitive strains (condition 2). Pooled plasmids from different conditions are used as template to amplify the gene of interest with primers having condition specific tags. Pooled PCR products are subject to deep sequencing (MiSeq Illumina platform) and the data is analyzed to get the relative ratio of the mutants in different conditions. (B) *In-vivo* activity assay for individual mutants in the CcdA library. Single mutants of CcdA were isolated, sequenced and individually transformed into the *E.coli* sensitive strain Top10. 5µl of the transformation mix was streaked onto LBamp agar plates and inoculated overnight at 37˚C to monitor the growth of mutants. Depletion ratio for each of the mutant spotted has been indicated in the table on left.

A

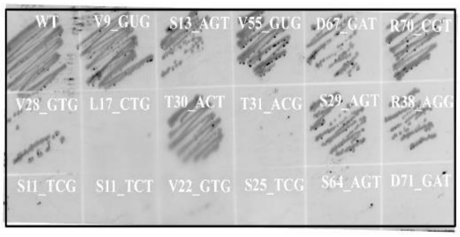


| | | | | | |
|---|---|---|---|---|---|
| 0.8 | 1.6 | 0.7 | 1.9 | 2 | 6.6 |
| 0.7 | 0.2 | 2.8 | 0.3 | 0.8 | 1.8 |
| 0.3 | 0.3 | 0.3 | 0.3 | 0.3 | 0.2 |

B

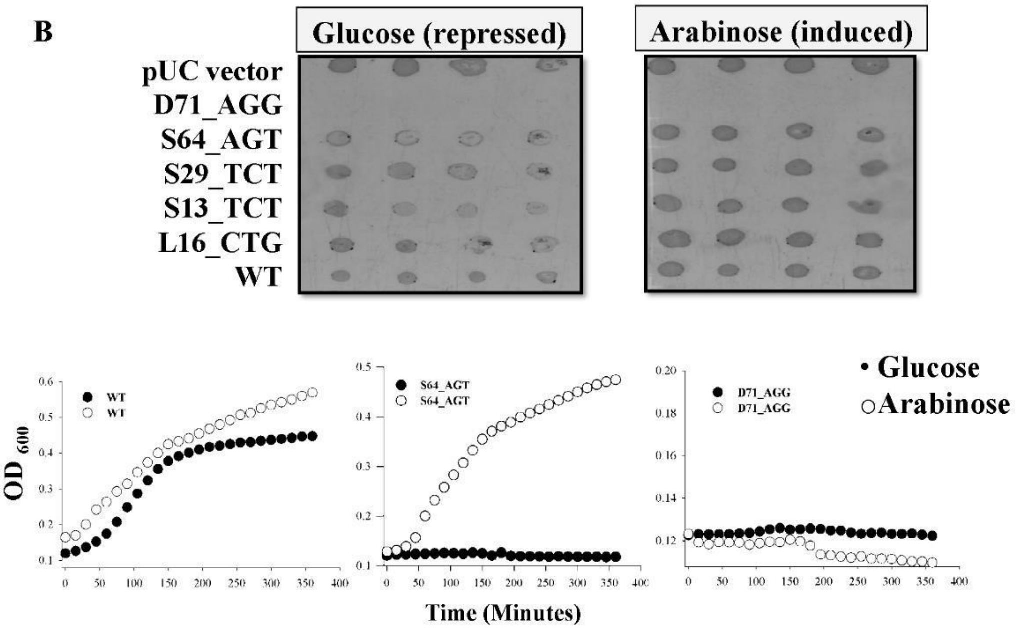


**Figure S2**: ***In-vivo* activity assay for individual synonymous mutants in the CcdA library.** (A) Single synonymous mutants of CcdA were isolated, sequenced and individually transformed into the *E. coli* sensitive strain Top10. 5µl of the transformation mix was streaked onto LBamp agar plates and inoculated overnight at 37˚C to monitor the growth of mutants. Depletion ratio for each mutant spotted has been indicated in the table on left. (B) Growth of single synonymous mutants in CcdA cloned under pBAD promoter in repressed vs induced conditions in Top10 strain. Except for D71R control and S64 synonymous mutants, the other synonymous mutants showed similar growth rates in resistant and sensitive strains but differed in their growth rates under inducing and repressed conditions.

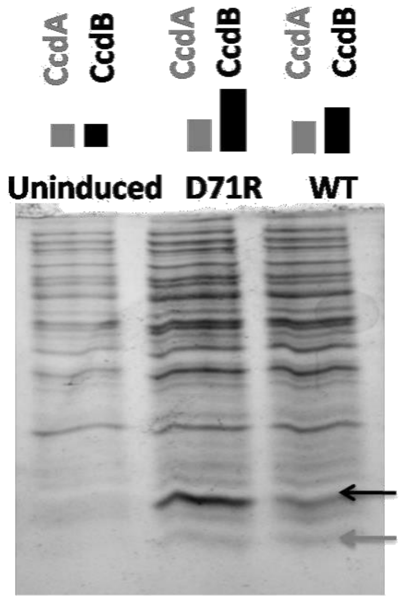


**Figure S3: Changes in the relative levels of the CcdA and CcdB proteins in the D71R mutant with respect to the WT monitored through SDS-PAGE.** The ccdAB operon was cloned under the pBAD, arabinose inducible promoter in the pBAD24 vector. The construct was transformed in the *E. coli* Top10Gyr cells, induced with 0.2% arabinose at an $OD_{600}$ of 0.6 and grown for 5 hours at 37°C. Cells were centrifuged, and the supernatant was subjected to 15% Tricine SDS-PAGE. Uninduced cells were used as a control and purified CcdA (grey) and CcdB (black) as molecular weight markers.

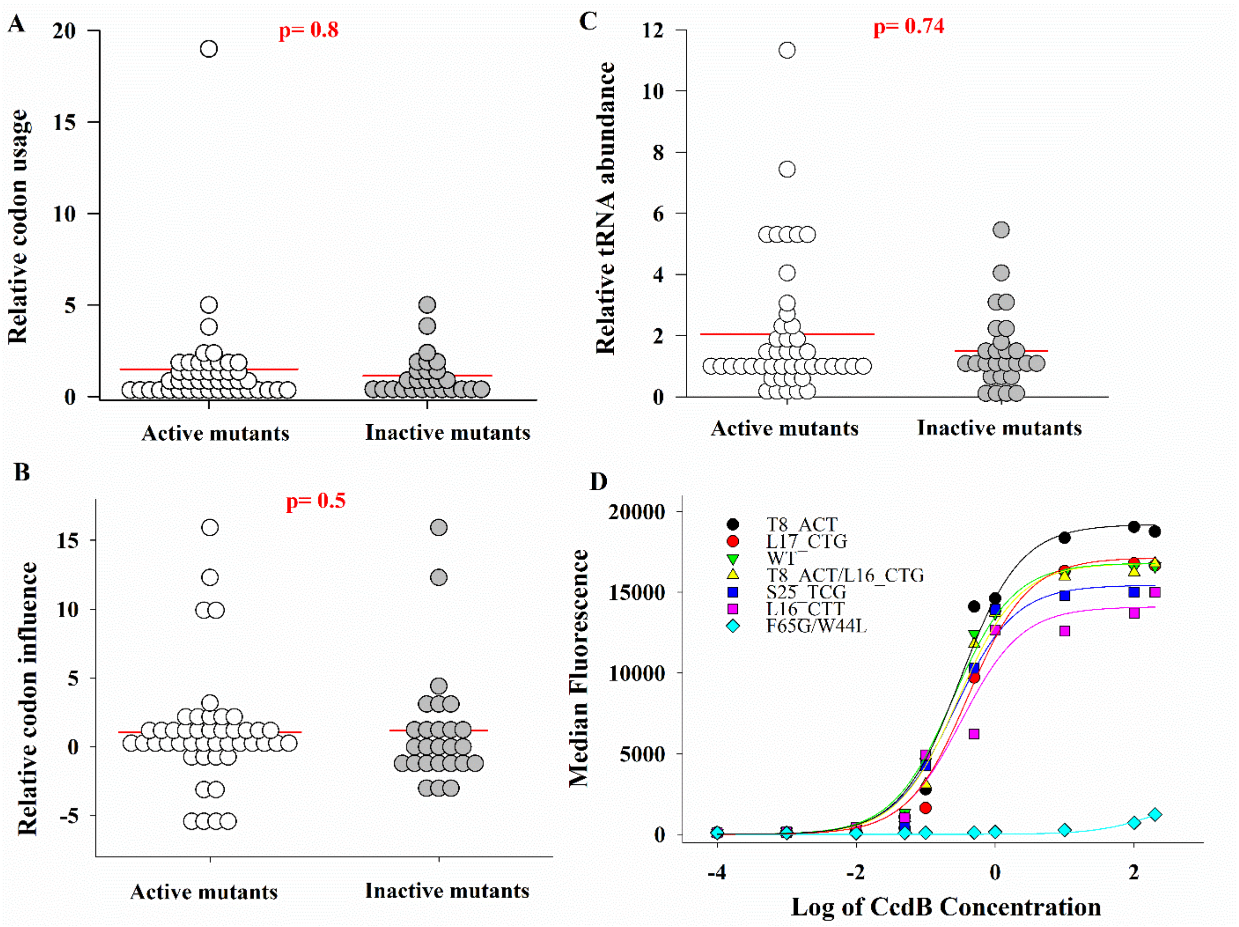


**Figure S4: Phenotype of CcdA synonymous mutants are uncorrelated with codon preference as well as CcdB binding.** (A-C) Single synonymous mutants in CcdA (N=71) were divided into active (N=45) and inactive (N=26) based on a depletion ratio cut-off of 0.3. Mutant distribution was analyzed with respect to codon usage (1), codon influence (2) and tRNA abundance (3) relative to the WT codon. (D) Assessing folding defects through binding of CcdA mutants to WT CcdB probed by yeast surface display. CcdA synonymous mutants were displayed on the yeast cell surface. CcdB binding was assessed by incubating cells with various concentrations of biotinylated CcdB, followed by binding to streptavidin conjugated AlexaFluor-633. Labelled cells were analyzed on a BD FACSARIA IIII and median fluorescence was calculated. All single synonymous mutants of CcdA showed comparable binding to WT. The double mutant F65G/W44L was taken as a negative control.

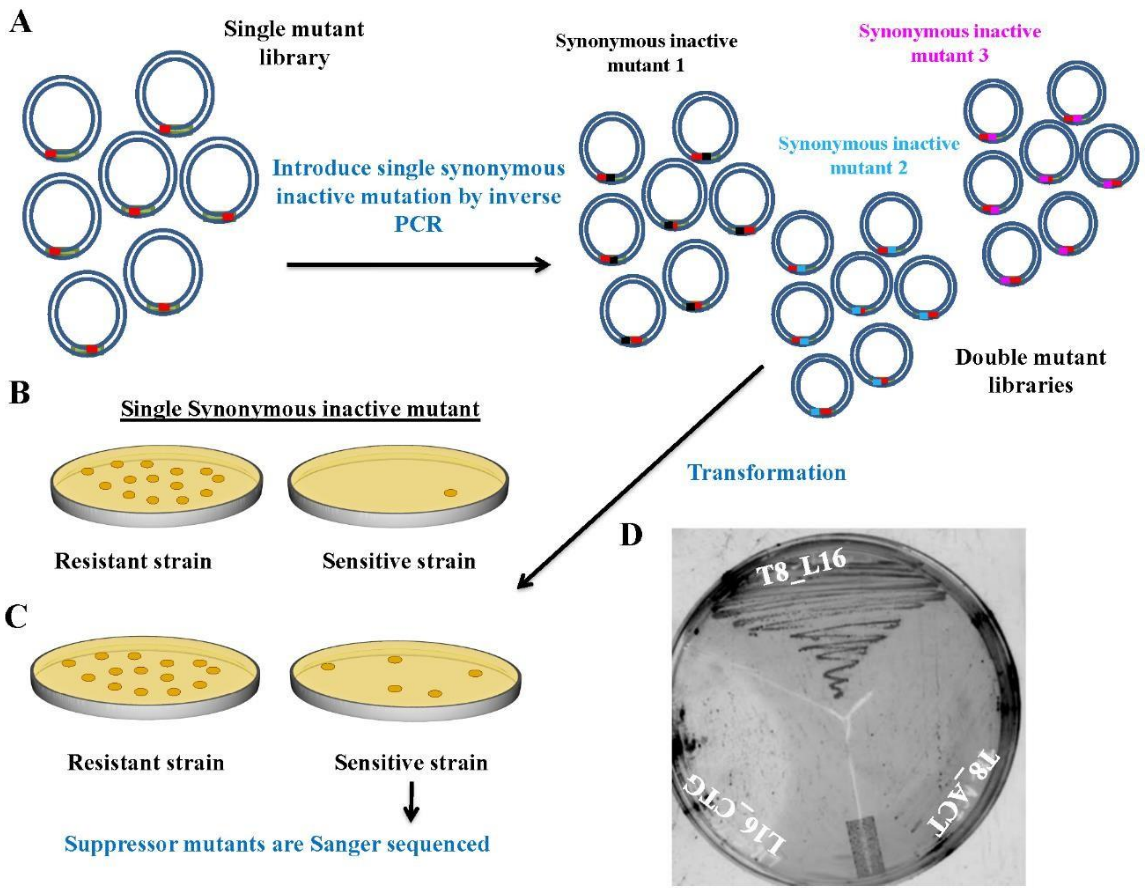


**Figure S5: Exhaustive suppressor analysis for synonymous inactive mutants.** (A) Different single synonymous, inactive mutations were introduced in the CcdA single site saturation mutagenesis library to create a double mutant library. (B) The single synonymous, inactive mutation leads to cell death the in sensitive strain. (C) Suppressor mutants rescue the inactive phenotype for the single synonymous mutation and can grow in the sensitive strain. These are identified by sequencing the entire operon. (D) Growth of Single mutants T8_ACT and L16_CTG and isolated suppressor double mutant T8_L16 transformed in *E.coli* Top10 sensitive strain in LBamp media.

GGAGATCCGAAAACCCCAAGTTACGGATCTTCCTCTCCCTCCGCACAGCGTTACATCCCGTCAGCACAGCATGTAGTGCCTCATACAGTTGCCCATGGCACTATAT

GTTGTGTTGTATCTCTGGACTGTGATGCGCCGCGCAGGGGCGGAAAACAGCGATATGATGATTTTCTCAGCGTTGTACACTTCCGGAAAGTCGTTTATTCAAATA

EcoRI

AAGTCGAATTCATACGAAACGGGAATGCGGTAATTACGCTTTGTTTTTATAAGTCAGATTTTAATTTTTATTGGTTAACATAACGAAAGGTAAAATACATAAGGCT

PsiI                                                                                Bgl2

TATAAAAAGCCAGATAACAGTATGCGTATTTGCGCGCTGATTTTTGCGGTATAAGAATATATACTGATATGTATACCCGAAGTATGTCAAAAAGATCTGTGCTATGA

AGCAGCGTATTACAGTGACAGTTGACAGCGACAGCTATCAGTTGCTCAAGGCGTATGATGTCAATATCTCCGGTCTGGTAAGCACAACCATGCAGAATGAAGC

CCGTCGTCTGCGTGCCGAACGCTGGAAAGCGGAAAATCAGGAAGGGATGGCTGAGGTCGCCCGGTTTATTGAAATGAACGGCTCTTTTGCTGACGAGAAC

DraI

AGGGACTGGTGAAATGCAGTTTAAAGTTTACACCTATAAAAGAGAGAGCCGTTATCGTCTGTTTGTGGATGTACAGAGTGATATTATTGACACGCCCGGGCG

ACGGATGGTGATCCCCCTGGCCAGTGCACGTCTGCTGTCAGATAAAGTCTCCCGTGAACTTTACCCGGTGGTGCATATCGGGGATGAAAGCTGGCGCATGATG

ACCACCGATATGGCCAGTGTGCCGGTCTCCGTTATCGGGGAAGAAGTGGCTGATCTCAGCCACCGCGAAAATGACATCAAAAACGCCATTAACCTGATGTTCT

BamHI

GGGGAATATAAATGTGAGGATCCGTTATACAC

**Figure S6**: Sequence of ccd operon synthesized and cloned in pUC57 vector at Genscript resulting in pUCccd plasmid. Restriction sites have been underlined and marked. The identified transcription start site is shown with an arrow. ccdA coding region is in green and ccdB is in grey. Mutations to facilitate cloning of CcdA and CcdB libraries are marked in red. Following the ccdB gene, the first few residues of the resD gene has been included with a stop at the second codon and mutation to introduce BamH1 site.
